# Development of iPSC-Derived T Cells Targeting EGFR Neoantigens in Non-Small Cell Lung Cancer

**DOI:** 10.1101/2025.02.15.638388

**Authors:** Kouta Niizuma, Toshinobu Nishimura, Jonathan Villanueva, Laura Amaya, Jonas L. Fowler, Taichi Isobe, Yusuke Nakauchi, Brandon Saavedra, Haojun Xu, Mahito Nakanishi, Adam C. Wilkinson, Kyle M. Loh, Joseph B. Shrager, Hiromitsu Nakauchi

## Abstract

A long-sought goal of cancer immunotherapy is to mass-produce T cells that specifically target tumor neoantigens. One decisive challenge is the identification of neoantigens derived from cancer driver genes. Here, we identify T cells that recognize the NSCLC-associated EGFR C797S mutation, which confers resistance to current inhibitors and is linked to poor prognosis. To overcome limitations in T cell availability, we reprogrammed EGFR C797S-specific T cells into induced pluripotent stem cells (iPSCs) and re-differentiated them into CD8 T cells. These iPSC-derived T cells specifically recognized the EGFR C797S mutation and effectively killed cancer cells expressing this mutation. Our findings underscore the potential of targeting driver mutation-derived neoantigens for immunotherapy and demonstrate that iPSC-derived T cells can mediate antitumor effects. Collectively, this approach combining neoantigen identification with T cell reprogramming may offer a promising strategy for targeting drug-resistant tumors.

## INTRODUCTION

Infusion of antigen-specific T cells into patients has led to unprecedented successes in treating certain types of cancer and inaugurated a new era in cancer therapy; however, substantial challenges remain. Two major challenges include (1) the scarcity of cancer-specific neoantigens to target and (2) the limited availability of T cells. In this study, we address both challenges in the context of lung cancer by integrating stem cell biology with immunological approaches.

The first challenge is to discover cancer-specific neoantigens^1^. While it remains controversial whether cancer-specific neoantigens exist for every tumor^2^, by definition mutations in cancer driver genes could hypothetically constitute cancer-specific neoantigens. This is exemplified by the recent finding that the mutant KRAS^G12D^ protein yields a neoepitope presented by HLA-C*08:02, which can be targeted by T cells^3,4^. Targeting neoantigens derived from driver (as opposed to passenger) genes would be preferable, as they would ensure that cancer cells bearing such mutations are eliminated. To overcome this challenge, here we describe a new pipeline to screen for cancer-specific neoantigens.

A second challenge is to create large numbers of T cells that recognize cancer-specific neoantigens, a challenge that is uniquely addressed by the advent of induced pluripotent stem cell (iPSC) technology. Themeli et al. demonstrated the proof of concept for mass production of CAR-T cells from CAR-transduced iPSCs^5^. Aside from CAR technology, we and others have demonstrated that small numbers of “exhausted” antigen-specific T cells can be isolated, reprogrammed into iPSCs, expanded in large numbers, and then re-differentiated to generate large amounts of antigen-specific rejuvenated T cells^6–9^. The repeatable and indefinite production of rejuvenated T cells is feasible by exploiting the self-renewal capacity of iPSCs^10^. Yet, this approach is still hampered by the scarceness of target cancer-specific neoantigens.

Here, we address these challenges in the context of lung cancer, which is the leading cause of cancer-related death in both men and women, estimated at 20-25% of all cancer death worldwide^11,12^. Lung cancer is one of the most aggressive malignancies and has a five-year survival rate of just ∼18%, substantially lower than many other leading cancers^13^. In non-small-cell lung cancers (NSCLCs), activating mutations in epidermal growth factor receptor (EGFR) are frequent, initially usually the L858R point mutation or small in-frame deletions in exon 19^14^. Treatment with first-generation small-molecule tyrosine kinase inhibitors (TKIs) leads to emerging resistance mutations such as T790M (the most common resistance mutation, seen in ∼60% cases of first-generation TKI-resistant patients). EGFR^T790M^*-*mutant NSCLC can be treated by third-generation TKIs such as osimertinib (AZD9291) and rociletinib (CO-1686)^15^, but subsequently the resistance mutations such as C797S arise^16^. The EGFR^C797S^ mutation confers resistance to all currently approved TKI drugs and is therefore currently challenging to treat in the clinic.

We hypothesized that drug-resistance mutations in *EGFR*, though challenging by conventional drugs, might produce immunogenic neoantigens recognizable by T cells. Here, we describe our workflow to identify and validate neoantigens derived from the EGFR^C797S^ mutation, and our isolation of T cells (and T-cell receptor sequences) that recognize an EGFR^C797S^ neoantigen. By reprogramming EGFR^C797S^-specific T cells into iPSCs and re-differentiating them into functional CD8^+^ T cells, we further demonstrate the feasibility for the generation of large numbers of EGFR^C797S^ neoantigen-specific T cells. To our knowledge, this study represents a novel approach to target the otherwise-undruggable EGFR^C797S^ mutation and demonstrates that of iPSC-derived T cells can mediate antitumor effects against solid tumors by targeting a cancer neoantigens. We believe that our approach may be generally applicable to discover neoantigens derived from cancer driver genes and to manufacture iPSC-derived T cells that target such neoantigens.

## RESULTS

### In silico approach for screening EGFR neoantigen candidates

In order to identify immunogenic neoantigens, first we searched TKI-resistant mutant EGFR protein sequences for potential immunogenic peptides that they might produce. We focused on three EGFR point mutations (L858R, T790M, and C797S) and tested whether they might be displayed on common HLA haplotypes (HLA-A*01:01, HLA-A*02:01, and HLA-A*11:01). As CD8^+^ cytotoxic T lymphocytes (CTLs) normally recognize 8-11 amino acid peptides^17^, we queried 8- to 11-mer protein sequences containing the single amino acid conversions (EGFR L585R, T790M, and C797S) as initial candidates, leading to 342 possibilities (query peptide × HLA) (Figure S1A).

To narrow down the list of candidate EGFR-mutant peptides that might be presented on HLA, we used peptide-HLA binding prediction algorithms. Although ∼15 prediction algorithms are available, no single algorithm was powerful enough to select potential peptides alone^18^. Thus, we used three popular algorithms, BIMAS, IEDB, and NetMHC, to score each candidate peptide (Figure S1B and Table S1). Scores from any two algorithms were poorly correlated (data not shown). Therefore, we selected all peptides that received a high peptide-HLA score on any of the three algorithms. For the original 342 peptides derived from mutant EGFR proteins, 88 passed this *in silico* selection step (Figure S1C).

### *In vitro* assessment of peptide-HLA binding affinity and stability

Next, we experimentally tested whether these mutant EGFR-derived peptides could bind HLA in two separate assays, by measuring their short-term binding (affinity) and long-term stability (Figure S1D). The short-term binding affinity assay identified 11 mutant EGFR peptides had significant affinity for HLA proteins (Figure S1E-S1G). Of these 11 peptide/HLA complexes, the long-term-binding assay (Figure S1H-S1J) identified two peptides derived from EGFR^C797S^ that stably bound HLA-A*02:01. These two peptides were: CS9.3 (QLMPFGSLL, τ1/2 = 85.70 [h]) and CS11.6 (LMPFGSLLDYV, τ1/2 = 89.10 [h]). As peptide/HLA complexes with high affinity and/or stability usually have higher immunogenicity^19,20^, we selected these two EGFR^C797S^-derived peptides, CS9.3 and CS11.6, as the top candidates for further analysis. We therefore synthesized these two peptides and tetramers for downstream functional T-cell assays, along with their counterpart WT peptides.

### Assessment of predicted epitopes in the context of immunogenicity

We sought to identify cytotoxic T lymphocytes (CTLs) and TCRs that recognized EGFR^C797S^-derived neopeptides CS9.3 or CS11.6 presented on HLA-A*02:01. We first screened CTLs obtained from the blood of HLA-A*02:01 human volunteers. From the same donors, we also obtained antigen-presenting cells (monocyte-derived dendritic cells [moDCs]^21^), which we pulsed with CS9.3 or CS11.6 peptides and then co-cultured with CD8^+^ CTLs isolated from the same donors to stimulate any peptide/HLA-specific CTLs that might exist within the repertoire of these human volunteers. After 2 weeks of coculture, we successfully identified tetramer^+^ CTL populations that bound to CS9.3 or CS11.6 (Figure 1A and S2A). These tetramer^+^ CTLs were isolated by fluorescence-activated cell sorting (FACS) and expanded again using Phytohemagglutinin-L (PHA-L) stimulation. These heterogenous CTL clones showed 60-90% reactivity specific to cognate tetramer and all displayed a terminal effector T cell phenotype (Figure 1B, 1C, S2B, and S2C).

**Figure 1:**
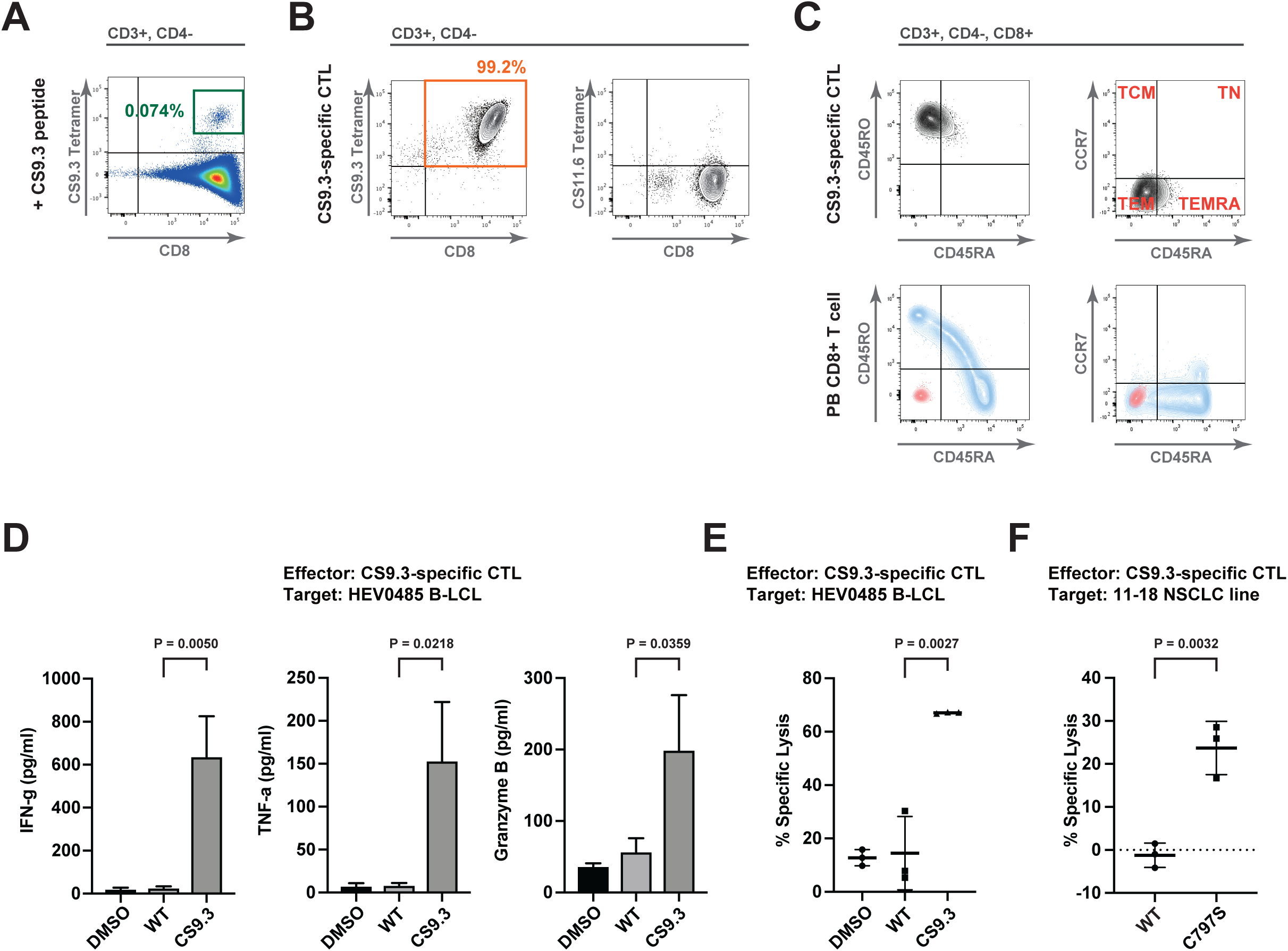
*In vitro* assessment of immunogenicity of EGFR neoantigens. (A) Flow cytometric analysis of the CD8^+^ T cells stimulated by peptide-pulsed autologous mDCs, 14 days after stimulation. Parent gate is indicated above each panel. (B) Flow cytometric analysis of purified and expanded CTL clones with cognate and irrelevant tetramers. Parent gate is indicated above each panel. (C) Flow cytometric analysis of memory phenotype in a CS9.3-specific CTL clone. Human peripheral blood CD8^+^ T cells (PB CD8^+^ T cell; bottom row) are shown as positive control (stained with the same antibody panel; blue) and negative control (unstained; red). Parent gate is indicated above each panel. (D) Cytometric Beads Array (CBA) analyses for IFN-γ, TNF-α, and Granzyme B secretion. CTL:HEV0485 = 10:1 (E:T ratio), 10 µg/ml peptide. (E) Antigen-specific cytotoxicity of CS9.3-specific CTL clone to HEV0485 B-LCLs. CTL:HEV0485 = 10:1 (E:T ratio), 10 µg/ml peptide. (F) Antigen-specific cytotoxicity of CS9.3-specific CTL clone to 11-18 NSCLC cell line. CTL:11-18 = 40:1 (E:T ratio). For (A-C), Data are representative of at least three independent experiments. For (D-F), Error bars = SD. Shown data are the representative in three times repeated experiments with technical triplicates in each experiment. P values are accorded to unpaired Student’s t test. See Figures S1 and S2 for additional data.

Next, we found that CS9.3- or CS11.6-specific CTL clones killed target cells expressing EGFR^C797S^-derived neopeptides, but not wild-type EGFR-derived peptides. To assess their antigen-specific cytotoxicity, CS9.3- or CS11.6-specific CTLs were mixed with antigen-presenting cells (HEV0485 B-LCL), which constitutively express HLA-A*02:01 and can be pre-loaded with wild-type or mutant peptide. Upon co-culture with mutant peptide-pulsed antigen-presenting cells, CS9.3- or CS11.6-specific CTLs became activated, secreting IFN-γ, TNF-α, and Granzyme B into the culture medium (Figure 1D and S2D). This response was not elicited by wild-type peptide. More significantly, CS9.3- or CS11.6-specific CTLs killed CS9.3- or CS11.6-pulsed HEV0485 B-LCL cells^22^, whereas minimal cytotoxicity was observed against controls (Figure 1E and S2E). In summary, both CS9.3- and CS11.6-specific CTL clones showed mutant peptide-specific activation and killing activity.

CS9.3- and CS11.6-specific CTL clones also demonstrated mutant peptide-specific cytotoxicity against NSCLC cells, more specifically the 11-18 NSCLC cell line expressing HLA-A*02:01^23^. As the 11-18 cell line expresses wild-type EGFR, we generated an EGFR^C797S^ expressing 11-18 cell line, termed 11-18^C797S^, by retroviral transduction. The CS9.3-specific CTL clones specifically and potently killed 11-18^C797S^ target cells, but not the parental (wild-type EGFR) 11-18 cells (Figure 1F). The CS11.6-specific CTL clones also showed killing activity, but also displayed some cross-reactivity to wild-type EGFR (Figure S2F). Based on these results, we prioritized CS9.3 for subsequent experiments. These results confirmed that naturally-processed CS9.3 and CS11.6 peptides (i.e. mutant EGFR^C797S^ proteins that were cleaved by the proteasome into CS9.3 and CS11.6 peptides and transported to the cell surface through the endoplasmic reticulum) are immunogenic and represent cancer-related neoantigens.

### Assessment of EGFR neoantigen-specific TCRs

To further characterize our neoantigen-specific CTLs, we sequenced the *TCRA* and *TCRB* genes in CS9.3- or CS11.6-specific CTL clones. Interestingly, the diversity of the *TCRA* and *TCRB* alleles in all clones was low (Figure S3A and S3B). The most frequent *TCRA* and *TCRB* gene sequences were selected (Figure S3C) and then cloned into a lentiviral expression vector carrying a truncated CD19 surface marker tag, to enable us to track transduced cells (Figure 2A). To initially assess whether these TCRs were functional, Jurkat T-cell lymphoma cells carrying T-cell activation fluorescent reporters (*NFAT:eGFP* and *NF-kB:eCFP*) were transduced with the CS9.3- or CS11.6-specific TCRs. TCR-engineered Jurkat reporter cells bound HLA tetramers carrying either the CS9.3 or CS11.6 neoantigens (Figure 2B and S4A).

**Figure 2:**
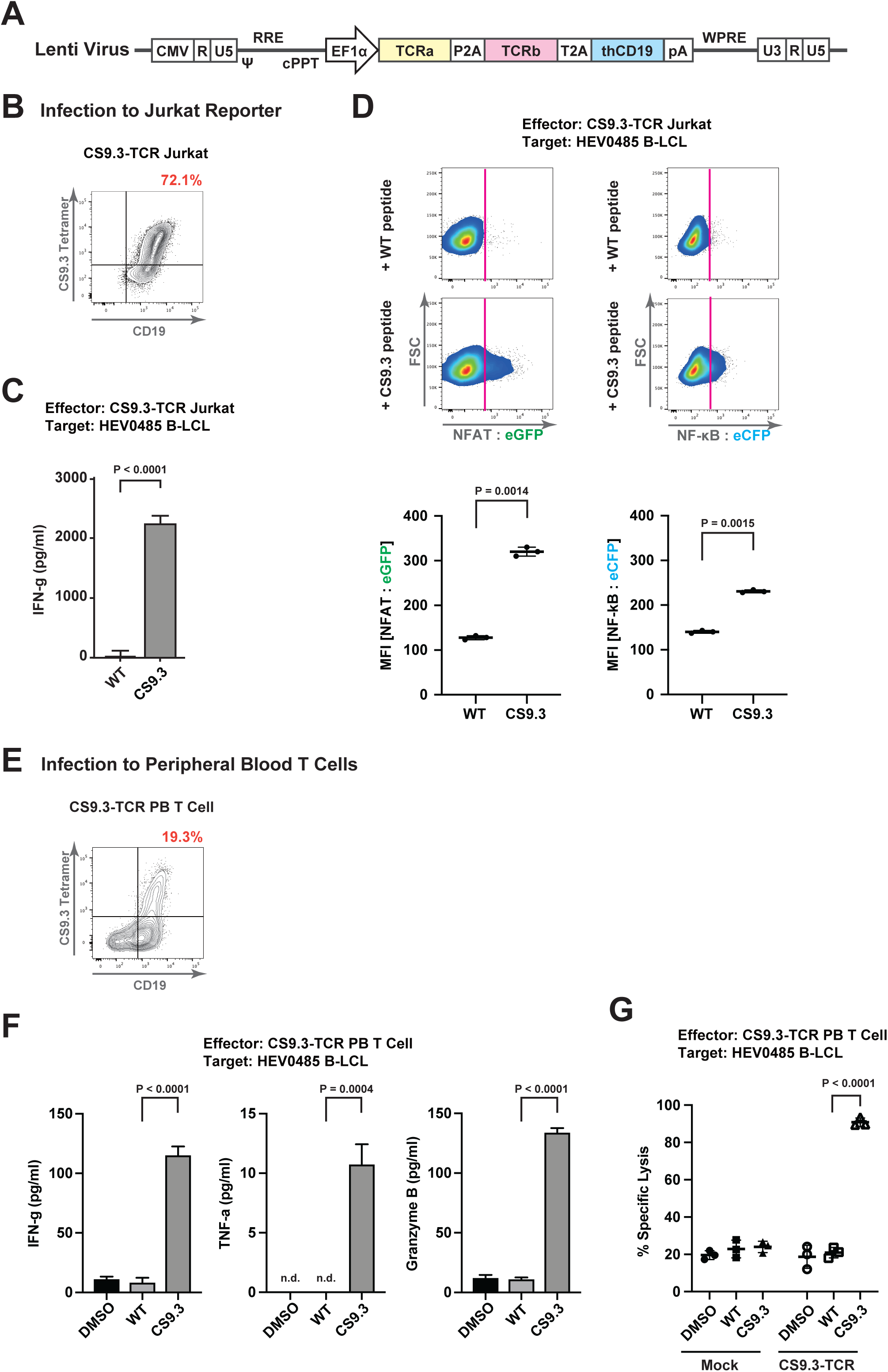
Functional TCR transfer and assessment of anti-cancer T cell activity. (A) Schematic illustration of lentivirus for TCR transfer. (B) Flow cytometric analysis for tetramer recognition by using Jurkat reporter cells infected by lentivirus encoding CS9.3-specific TCRs. (C) Activation and secretion of IFN-γ in CS9.3-TCR transferred Jurkat reporter cells. Jurkat:HEV0485 = 10:1 (E:T ratio), 10 µg/ml peptide. (D) Flow cytometric analysis of CS9.3-TCR transferred Jurkat reporter cells. Jurkat:HEV0485 = 10:1 (E:T ratio), 10 µg/ml peptide. Bottom panels show the mean of fluorescence intensity (MFI) of reporters. (E) Flow cytometric analysis for tetramer recognition by using peripheral blood CD8^+^ CTLs infected by lentivirus encoding CS9.3-specific TCRs. (F) Cytometric Beads Array (CBA) analyses for IFN-γ, TNF-α, and Granzyme B secretion in CS9.3-TCR transferred CTLs. CTL:HEV0485 = 10:1 (E:T ratio), 10 µg/ml peptide. (G) Antigen-specific cytotoxicity of a CS9.3-TCR transferred CTLs to HEV0485 B-LCLs. CTL:HEV0485 = 10:1 (E:T ratio), 10 µg/ml peptide. Effector T cells shown here had ∼30% tetramer^+^. For (C, D, F, and G), Error bars = SD. Shown data are the representative in three times repeated experiments with technical triplicates in each experiment. P values are accorded to unpaired Student’s t test. See Figures S3 and S4 for additional data.

To confirm that these CS9.3- or CS11.6-specific TCRs confer cytotoxic responses against these neoantigens, we co-cultured TCR-engineered Jurkat reporter cells with target cells (HEV0485 B-LCL) that were pulsed with CS9.3 or CS11.6 peptides^24^. When the CS9.3 neoantigen was presented to CS9.3-TCR-expressing Jurkat reporter cells by HEV0485 B-LCL target cells, the Jurkat reporter cells became activated and secreted IFN-γ (Figure 2C), and the NFAT and NFkB T-cell activation reporters were also triggered (Figure 2D). Although CS11.6-TCR-expressing Jurkat reporter cells secreted IFN-γ, no reporter activity was observed (Figure S4B and S4C). Various reasons may explain this lack of activity, including inefficiency of TCR transduction, instability of the TCR complex, and/or incompatibility with endogenous TCRs.

Finally, to assess the activity of the TCRs in primary human T cells, peripheral blood CD8^+^ CTLs were transduced with the CS9.3-TCR or CS11.6-TCR lentivirus. As CS11.6-TCR again displayed no reactivity to the tetramer, it was removed from further analysis (Figure S4D). By contrast, primary human T cells engineered with the CS9.3-TCR CTLs potently bound to CS9.3:HLA-A*02:01 tetramer (Figure 2E). Additionally, cytokine secretion and neoantigen-specific killing were observed for CS9.3-TCR expressing CTLs (Figure 2F and 2G); responses against wild-type EGFR were not detected. In summary, our data clearly demonstrate that the cloned CS9.3-TCR has neoantigen-specific avidity and confers cytotoxic functions to transduced CTLs. We therefore conclude that the CS9.3 peptide derived from EGFR^C797S^ proteins is a highly-potent immunogenic neoantigen.

### Generation of neoantigen-specific CTLs via iPS-reprogramming and redifferentiation

We and others have previously reported the efficient reprogramming of antigen-specific CTLs into iPSCs and their subsequent redifferentiation into CTLs from a small initial population, addressing the critical limitation of limited T cell numbers available for adoptive immunotherapy^6–9,25^. As there are no current TKI treatments for EGFR^C797S^-mutant cancers, we wondered whether it would be possible to generate CS9.3-specific iPSC-derived CTLs (iCTLs) in quantities sufficient for therapeutic applications.

To generate large numbers of iCTLs (Figure 3A), first we reprogrammed CS9.3-specific CTLs into “T-iPSCs” (i.e., iPSCs that carry rearranged *TCR* genes) using Sendai virus (expressing OCT3/4, SOX2, KLF4, c-MYC, and SV40 large T antigen)^9^. We then re-differentiated these CS9.3-specific T-iPSCs into CTLs. Specifically, CS9.3-specific T-iPSCs were differentiated into hematopoietic progenitor cells^26^, followed by differentiation into CD4/CD8 double positive T-lineage cells in artificial thymic organoid (ATO) cultures by mixture with NOTCH-ligand-expressing stromal cells^27,28^ (Figure S5A), and then CD4/CD8 double positive cells were further matured into CD8 single positive CTLs (iCTLs) with stimulation by PHA-L (Figure 3B). Transcriptional fidelity of T cell differentiation was further confirmed using a gene set of differentially immune cell genes, with iPSC-derived CTL clones had displaying a gene expression profile similar to primary CD8^+^ T cells (Figure 3C).

**Figure 3:**
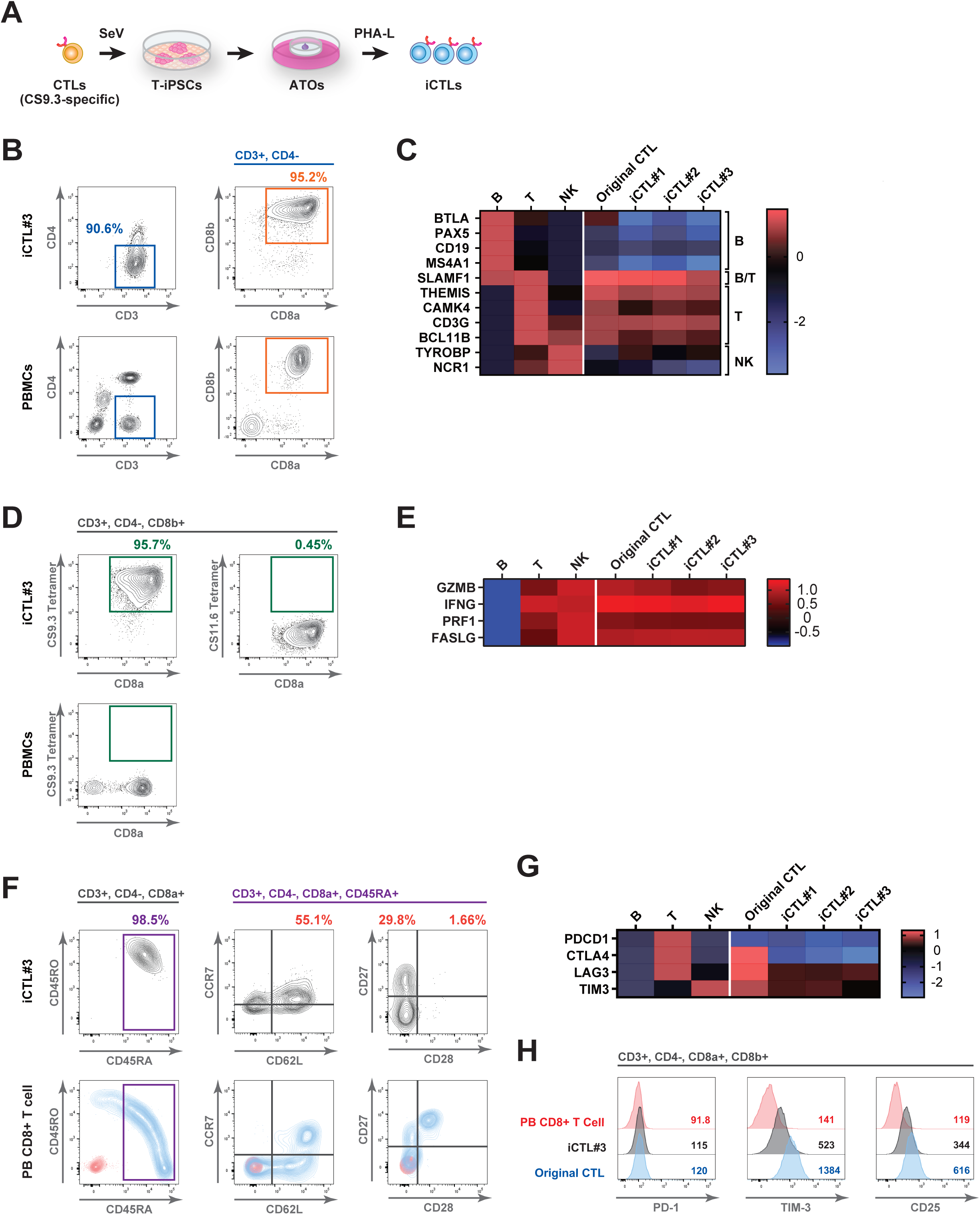
Generation of iCTLs via iPS-reprogramming and redifferentiation. (A) Schematic illustration of the generation of iCTLs. (B) Flow cytometric analysis of T cell markers in iCTL#2 clone. Human peripheral blood mononuclear cells (PBMCs; bottom row) are shown as positive and negative controls. Parent gate is indicated above each panel. (C) qPCR analysis for B-cell, T-cell, and NK-cell markers. Heatmap coloring reflects the relative expression of the genes using a log2 scale standardized by the expression of YWHAH. Peripheral blood B cells, T cells, and NK cells were analyzed in parallel as positive control and negative control of expression. (D) Flow cytometric analysis of antigen-specificity in iCTL#2 clone with cognate (CS9.3) and irrelevant (CS11.6) tetramers. Human peripheral blood mononuclear cells (PBMCs; bottom row) are shown as negative controls. Parent gate is indicated above each panel. (E) qPCR analysis for cytotoxic molecules. Heatmap coloring reflects the relative expression of the genes using a log2 scale standardized by the expression of YWHAH. Peripheral blood B cells, T cells, and NK cells were analyzed in parallel as positive control and negative control of expression. (F) Flow cytometric analysis of memory phenotype in iCTL#2 clone. Human peripheral blood CD8^+^ T cells (PB CD8^+^ T cell; bottom row) are shown as positive control (stained with the same antibody panel; blue) and negative control (unstained; red). Parent gate is indicated above each panel. (G) qPCR analysis for T-cell exhaustion markers. Heatmap coloring reflects the relative expression of the genes using a log2 scale standardized by the expression of YWHAH. Peripheral blood B cells, T cells, and NK cells were analyzed in parallel as positive control and negative control of expression. (H) Flow cytometric analysis of T-cell exhaustion markers in iCTL#3 clone. Mean of fluorescence intensity (MFI) of each sample was shown. Human peripheral blood CD8^+^ T cells (PB CD8^+^ T cell) are shown as negative control. Parent gate is indicated above each panel. See Figures S3 for additional data.

Upon differentiation, iCTLs had the same antigen-specificity as original CS9.3-specific CTLs, as they bound CS9.3:HLA-A*02:01 tetramer (Figure 3D). Such CTLs also demonstrated cytotoxic molecules expression (Figure 3E) and displayed a rejuvenated naïve/memory immunophenotype (Figure 3F). Down-regulation of T-cell exhaustion markers was observed by qPCR and flow cytometry (Figure 3G and 3H). However, as PD-1 expression was low in an original CS9.3-specific CTL clone, there was no significant difference of the PD-1 expression before and after iPSC-reprogramming.

We next demonstrated that CS9.3-specific iCTLs were functional: they specifically killed target cells (HEV0485 B-LCL) that were pulsed with CS9.3, but not wild-type EGFR-derived, peptides (Figure 4A). Similar results were obtained when iCTLs were mixed with K562 aAPC expressing single chain trimer (peptide-B2M-HLA-A*02:01) or 11-18^C797S^ NSCLC cell line (Figure 4B and 4C). Consistent with the T cell profile of iCTLs (Figure 3C), we further confirmed that the iCTLs did not have NK cell-like, antigen-independent cytotoxicity by co-cultureing iCTLs clones with K562 (Figure S6). These results therefore confirm that iCTLs demonstrate neoantigen-specific cytotoxic activity. Since T-iPSCs are capable of expanding infinitely, our approach enables an unlimited supply of iCTLs to be made for repeated administration into patients^10^. This therefore opens the prospect of treating EGFR^C797S^-mutant NSCLC, as this mutant EGFR protein cannot be currently targeted by TKIs but we find that it yields a cancer-specific neoantigen that can be targeted by iPSC-derived rejCTLs.

**Figure 4:**
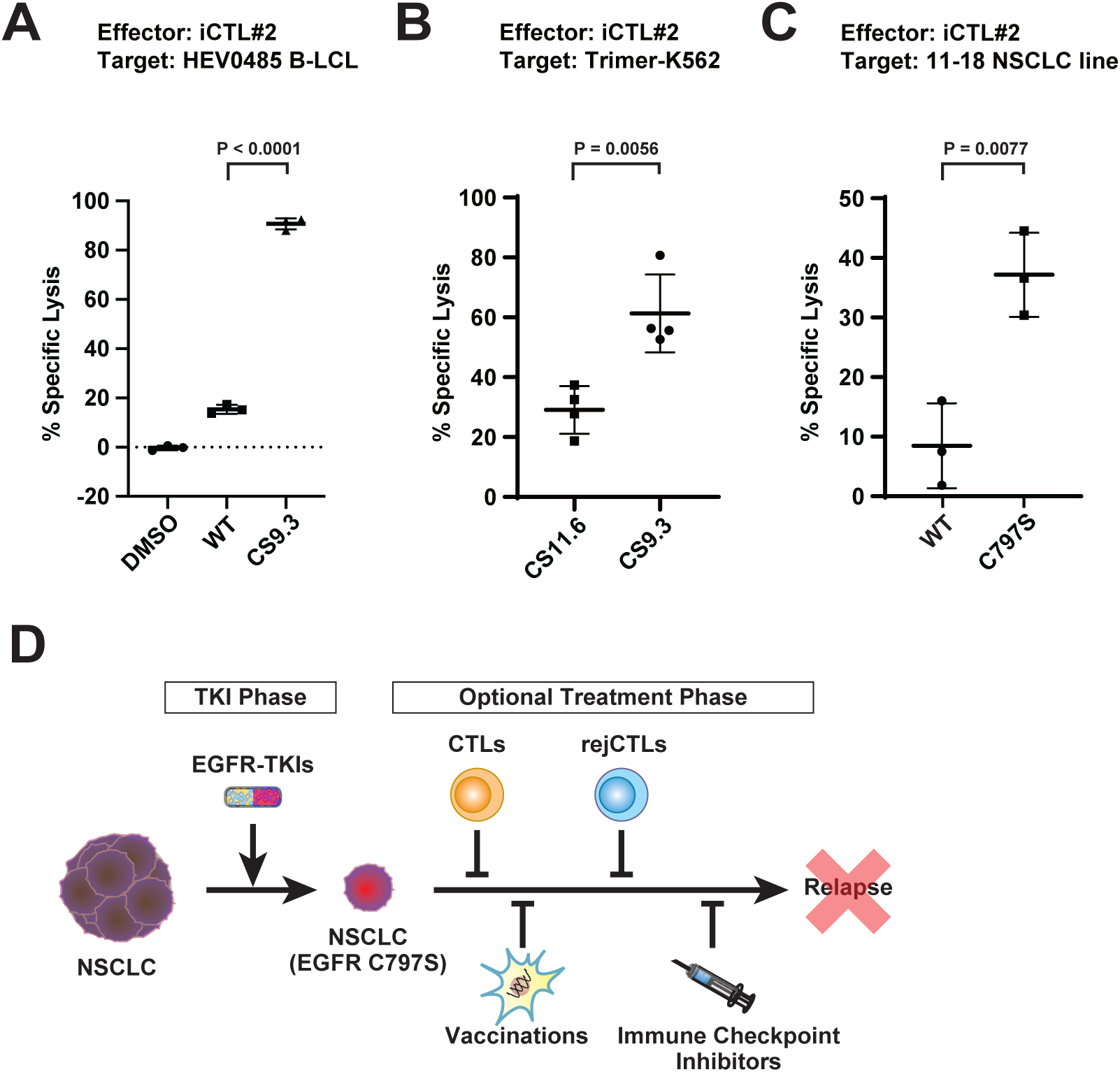
Revitalized antigen-specific cytotoxicity of rejCTLs. (A) Antigen-specific cytotoxicity of iCTL#2 clone to HEV0485 B-LCLs. CTL:HEV0485 = 10:1 (E:T ratio), 10 µg/ml peptide. (B) Antigen-specific cytotoxicity of iCTL#2 clone to Trimer-K562. CTL:Trimer-K562 = 2.5:1 (E:T ratio). (C) Antigen-specific cytotoxicity of iCTL#2 clone to 11-18 NSCLC cell line. CTL:11-18 = 20:1 (E:T ratio). (D) Schematics of combination immunotherapy of EGFR-TKI with optional treatments (CTLs, repeated administration of iCTLs, vaccinations, and immune checkpoint inhibitors). For (A-C), Error bars = SD. Shown data are the representative in three times repeated experiments with technical triplicates or quadruplicate in each experiment. P values are accorded to unpaired Student’s t test.

Since T-iPSCs are capable of expanding infinitely, and are capable of accepting multiple gene edits to convert them into hypoimmunogenic iPSCs^29–31^, our approach may enable an unlimited supply of rejuvenated CTLs to be made available as an allogeneic “off-the-shelf” therapy for a broader range of NSCLC patients carrying the EGFR^C797S^ mutation ^10^(Figure 4D). Our work therefore opens the prospect of treating EGFR^C797S^-mutant NSCLC, a cancer type that cannot be currently targeted by TKIs but one that we have found yields a cancer-specific neoantigen targetable by iPSC-derived rejuvenated CTLs.

## DISCUSSION

Here, we address two challenges to the use of antigen-specific T cells for cancer therapy: (1) the scarcity of cancer-specific neoantigens to target and (2) the limited availability of T cells. Although considerable effort has been devoted to identifying cancer-specific neoantigens, those derived from driver genes remain rare. Notable examples include a mutant KRAS^G12D^ protein-derived neoepitope presented by HLA-C*08:02^3,4^ and TP53^R175H^ protein-derived neoepitope presented by HLA-A*02:01^32^. Furthermore, a recent study targeting neoantigens derived from EGFR^T790M/C797S^ mutations in non-small cell lung cancer has demonstrated the potential of peptide-based immunotherapy in this setting^33^; however, that work was limited to short-term binding affinity assays and peptide vaccination strategies.

In our study, we identified CS9.3, a neoantigen derived from the EGFR C797S mutant protein, as one of the few cancer-specific neoantigens originating from a driver gene. Moreover, because CS9.3 is presented on HLA-A*02:01—the most frequent HLA haplotype in the human population—our approach has broad potential applicability. In addition, the cross-reactivity of high-affinity peptides to other HLA-A02 subtypes^34^ may further extend the benefits of targeting CS9.3 in NSCLC. Given that EGFR^C797S^ mutation arises in a NSCLC driver gene and is unlikely to be present in normal tissues, targeting CS9.3 may help to minimize the off-target toxicities often encountered with T cell therapies that target antigens expressed in both cancerous and healthy cells.

Regarding the second challenge, the establishment of neoantigen-specific T-iPSC lines has the potential to greatly impact cancer immunotherapy, opening up a new venue to “off-the-shelf” adoptive T cell immunotherapy. Along with the retained antigen-specificity and functional characteristics, T-iPSC-derived rejuvenated CTLs (rejCTLs) offer additional benefits by enabling the scalable production and the banking of functionally validated CTLs. The endless supply of neoantigen-specific rejCTLs enables us to treat cancer patients, overcoming the limitations in conventional T cell immunotherapies, which are constrained by the limited cell numbers obtainable through *ex vivo* expansion and the exhaustion associated with repeated stimulation. Our CS9.3-specific rejCTLs demonstrated the retention of TCR expression, robust *in vitro* cytotoxicity against CS9.3-expressing cell lines. Importantly, these rejCTLs are neoantigen-specific, as they do not target wild-type EGFR. The monoclonality of the TCRs in the rejCTL population also reduces the risk of off-target toxicities when the TCR has been well-characterized prior to manufacturing. Accordingly, further development of this candidate will require an in-depth search for cross-reactivity against the normal human peptidome, as has become standard in the field of TCR based therapeutics^35^.

Recent advancement in cancer immunotherapy including immune checkpoint blockade and adoptive transfer of antitumor lymphocytes, which have demonstrated the power of cytotoxic T cells in treating various types of solid tumors. *Ex vivo*-expanded tumor-infiltrating lymphocytes (TILs) have shown efficacy in some melanoma patients^36^. Target-redirection by transferring an antigen-specific T cell receptor (TCR) or chimeric antigen receptor (CAR) have shown dramatic effectiveness against tumors with specific target molecules via TCR or CAR recognition^37^. However, these types of T cell immunotherapy have mainly been conducted in autologous settings, and thus are costly, time-consuming, and depend on the quality of the patient’s T cells, and sometimes ends in failure to produce effector cells. We and others are developing a parellel method for the preparation of antitumor rejCTLs by incorporating iPSC technology into the development of T cell immunotherapies. T-iPSCs opens up a range of exciting possibilities for equipping rejCTLs with better features via gene editing, e.g. addition of activation molecules and/or removal of inhibition molecules^38^, camouflaging their HLAs and CD47 expression to avoid rejection by recipient’s immunity^29,39^, and/or introduction of a safety switch system (Nishimura et al., 2020; Martin et al., 2020). Therefore, T-iPSC technology offers a master cell banking of rejCTLs with a well-defined quality, and envision its use as an allogeneic “off-the-shelf” therapy for broad range of patients with the same cancer type or mutations.

Most mutations in cancerous cells are random passenger mutations that are not conserved across patients in the same tumor type. “Personalized” (or “private”) T cell therapy generally targets these private neoantigens^40^. On the other hand, T-iPSC system goes well together with “shared” (or “public”) neoantigens, which are generally the product of strong driver mutations responsible for the malignant transformation in many patients. Recurrent “public” neoantigens allows us to prepare neoantigen-specific T-iPSCs with extensive gene editing for the application in off-the-shelf therapies. Excitingly, cancer-specific “public” neoantigens such as CS9.3 are not restricted to the usage in T-cell immunotherapy, but may also find application in cancer vaccination or as biomarkers for immune checkpoint inhibitor therapy^41^. The identification of new neoantigens therefore has broad and important implications for cancer treatment.

While our current study demonstrates the potential of CS9.3-specific iPSC-derived CTLs through *in vitro* evaluations, future studies should aim to rigorously assess the *in vivo* efficacy, persistence, and safety of these cells in appropriate animal models to fully realize their therapeutic potential.

## EXPERIMENTAL PROCEDURES

### Regulatory and institutional review

All animal experiments were conducted according to experimental protocols approved by the Stanford Administrative Panel on Laboratory Animal Care (APLAC-28406) and in adherence with the US National Institutes of Health’s Guide for the Care and Use of Laboratory Animals. All human pluripotent stem cell experiments were conducted in accord with experimental protocols approved by the Stanford Stem Cell Research Oversight (SCRO) committee. All experiments using human patient specimens were conducted in accordance with a protocol approved by the Administrative Panel on Biosafety (APB). The entire study was conducted in accordance with the Declaration of Helsinki.

### Cell lines

K562 and K562 aAPC, including its derivative Trimer-K562, were grown in R10 medium (RPMI 1640 + 10% FBS + 1% GlutaMax), splitting 1:20 in every 3 days. MS5-hDLL1 cells were grown in R10 medium, splitting 1:10 in every 3 days. 11-18 cells were grown in R10 medium, splitting 1:20 in every 7 days.

### Flow cytometry

All sample staining for flow cytometry were performed in FACS buffer (PBS + 0.5% BSA + 2 mM EDTA + 0.09% sodium azide) for 30 min on ice. In case two or more antibodies conjugated with brilliant violet dye were used, Brilliant Stain Buffer was added to samples prior to antibody staining. For tetramer co-staining, tetramer was added to cells (1:50 final dilution) 20 min prior to additional antibody staining. Finally, washed cell pellets were resuspended in FACS buffer containing propidium iodide (PI) or 7-amino-actinomycin D (7-AAD) and were strained through a 40 µm filter. Analysis was performed on BD FACS Aria II instrument at Stanford Stem Cell Institute FACS Core.

### Isolation of peripheral blood mononuclear cells

Human white blood cells from were purchased from Stanford Blood Center and mononuclear cells were isolated by density gradient centrifugation using Ficoll-Paque Plus and SepMate™-50. Isolated leukocytes were washed twice in FACS buffer (PBS + 0.5% BSA + 2 mM EDTA + 0.09% sodium azide), then treated with ACK Lysing Buffer for 5 min at room temperature. HLA type of each leukocyte sample was checked by flow cytometer with fluorophore-conjugated anti-HLA-A1, -A2, or -A24 antibodies.

### HLA typing

Genomic DNA was extracted from 1-2 million PBMCs using a DNeasy Blood & Tissue Kits. HLA typing was performed to subtype-level (4-digit) resolution using TruSight HLA v2 Sequencing Panel. Next generation sequencing was conducted in Stanford Protein and Nucleic Acid Facility (PAN) with Illumina MiSeq at paired-end 2 x 150 bp depth. Data were analyzed by Assign 2.0 TruSight HLA Analysis Software.

### Generation of monocyte-derived DCs

Monocyte-derived DCs (moDCs) were induced by following previous report^21^. In brief, CD14^+^ cells were isolated by immunomagnetic selection using anti-CD14 microbeads and LS magnetic column. At this point, the purity of CD14^+^ cells were >95% in average. Cells were resuspended in DC medium (CellGenix^®^ GMP DC medium + 1% human AB serum) at 1 x 10^6^ cell/ml, then 2 ml was plated into each well of 6-well plate. For 2 days, cells were cultured in 3 ml DC medium with 10 ng/ml IL-4 and 86 ng/ml GM-CSF. At day 3, 1.5 ml of fresh DC media with 10 ng/ml IL-4 and 172 ng/ml GM-CSF was supplemented. At day 4, both floating and adhesive cells were collected by vigorous pipetting, pelleted, and resuspended in DC medium with 10 ng/ml IL-4, 86 ng/ml GM-CSF, 10 ng/ml LPS, 10 ng/ml IFN-g, and 2.5 ug/ml custom synthesized peptide at 1 x 10^6^ cells/ml. Cells were transferred in each well of 6-well plate (2 ml per well) and cultured for one day. All cells were then collected by vigorous pipetting for in T cell stimulation.

### Generation of neoantigen-specific cytotoxic T lymphocyte (CTL) clones

After the isolation of CD14^+^ cells, PBMCs of the same donor were processed for isolation of CD8^+^ CTLs by using CD8^+^ T Cell Isolation Kit and LS magnetic column. Purified CTLs (>95% purity) were resuspended in R10ITS medium (RPMI 1640 + 10% FBS + 1% GlutaMax + 1% ITS-X). Peptide-pulsed moDCs were treated by mitomycin C (MMC) at 37°C for 2 hours, then washed in DPBS twice. CTLs were mixed with MMC-treated moDCs (CTL:moDC = 3:1) and cultured in R10 medium with 10 ng/ml IL-7 and 10 ng/ml IL-15 for one week. After one week, a second stimulation was performed (as above) and cells were cultured for one more week. After two weeks, all cells were collected and stained with anti-human CD3, CD4, CD8a antibodies, and tetramer in FACS buffer at 4°C for 30 min. CTLs were analyzed by FACS Aria II and tetramer^+^ cells were purified.

Purified tetramer^+^ CTLs were expanded by Rapid Expansion Protocol (REP) with slight modification^42^. A total of 10 × 10^6^ cells and 10 × 10^6^ of Dynabeads™ Human T-Activator CD3/CD28 were mixed in a 1.5 ml Eppendorf tube, then rotated up and down for 45 min at room temperature. The 1.5 ml tube was then placed on a DynaMag™-2 Magnet for 2 min, and after the beads pelleted, the medium was aspirated completely. The beads/cells were resuspended in 10 ml of R10ITS medium (= 1.0 × 10^6^ cells/ml) supplemented with 10 ng/ml IL-7 and 10 ng/ml IL-15, and subsequently culture in 24-well plate wells at 1.0 × 10^6^ cells/ml/cm^2^.

### Generation of human induced pluripotent stem cells (iPSCs)

From CS9.3- or CS11.6-specific CTLs, iPSCs were generated by following the method in a previous report^9^. In brief, CTLs were stimulated by 5 ng/ml PHA-L in R10ITS medium (RPMI 1640 + 10% FBS + 1% GlutaMax + 1% ITS-X) for 2 days, then transduced with reprogramming factors via Sendai virus vectors. Transduced cells were seeded onto murine embryonic fibroblasts (MEF) feeder cells and cultured in myeloid medium, which was gradually replaced with conventional human iPSC medium (DMEM/F12 HAM + 20% knockout serum replacement + 1% GlutaMax + 1% nonessential amino acids + 50 µM 2-mercaptoethanol + and 5 ng/ml bFGF). The iPSC colonies >1 mm diameter were picked mechanically, dissociated into small clumps by pipetting in 1.5 ml Eppendorf tube, and transferred onto iMatrix-511 coated plate with 10 µM Y-27632 in StemFit® Basic02 medium supplemented with 20 ng/ml bFGF.

### Human pluripotent stem cell (PSC) culture and differentiation

CS9.3-specific T-iPSCs were routinely propagated feeder-free in StemFit® Basic02 medium supplemented with 20 ng/ml bFGF on cell culture plastics with iMatrix-511 basement membrane matrix, with single cell dissociation by using TrypLE™ Express Enzyme^43^. Undifferentiated hiPSCs were maintained at high quality with particular care to avoid any spontaneous differentiation, which would confound downstream differentiation. CS9.3-specific T-iPSCs were differentiated into hematopoietic progenitor cells using a non-transgene differentiation method as previously described^26^, followed by differentiation into CD4/CD8 double positive cells by artificial thymic organoid (ATO) culture^27,28^.

### Cytometric beads array (CBA)

The protein level of IFN-γ, TNF-α, and Granzyme B in the culture supernatants of cytotoxicity assay were measured by CBA. The culture supernatants were collected 24 hours after the stimulation and stored at -80 until the analysis. Samples were analyzed on the BD FACS Aria II Flow Cytometer using FCAP ArrayTM software according to the manufacturer’s instruction.

### *In vitro* cytotoxicity assay measured by bioluminescence

Firefly Luciferace (FLuc) was introduced into HEV0485 B-LCL, K562, Trimer-K562, and 11-18 cell line by a transposon system: 1 µg of Piggyback donor plasmid (pPB_CAG_FLuc_IRES_Neo) and 2 µg of transposase plasmid (pCMV-pBase) were co-transfected into 5 × 10^5^ cells by using Lipofectamine™ 3000 Transfection Reagent. Transfected cells were cultured in the presence of 200 ng/ml G418 to select stably FLuc expressing cells.

5 ×10^3^ FLuc-expressing 11-18 cells (target) were placed in black-wall 96–well flat bottom plates in triplicates and incubated overnight at 37°C, then checked the adhesion. 5 ×10^3^ FLuc-expressing HEV0485 B-LCL, K562, or Trimer-K562 cells (target) were placed in black-wall 96–well flat bottom plates in triplicates on the day of starting the assay. Subsequently, effector cells were added at various effector-to-target (E:T) ratios, more than 4 conditions, and incubated at 37°C for 20-24 hours. BLI was then measured for 10 seconds with IVIS Spectrum. Cells were treated with 1% Nonidet P-40 (NP40) as a measure of maximal killing control. Target cells incubated without effector cells were used to measure spontaneous death control. Triplicate wells were averaged and percent lysis was calculated from the data with the following equation: % specific lysis = 100 – (Flux[test] – Flux[maximal killing]) / (Flux[spontaneous death] – Flux[maximal killing])

×100.

### Sequencing of T-cell receptor CDR3 region

CDR3 sequences of rearranged TCRA and TCRB genes in CTL clones were determined by Amp2Seq service (iRepertoire, Inc.). Paired-end sequencing was performed on Illumina MiSeq, for an average read depth of 500,000 reads per sample. Whole nucleotide sequences of rearranged TCRA and TCRB were reconstructed by fetching nucleotide sequences of V-, D-, or J-segment from IMGT database, then translated into protein sequences. Codon optimized nucleotide sequences were produced by using the protein sequence and Codon Optimization Tool (IDT).

### mRNA expression analysis by qPCR

Total RNA was isolated using RNeasy Micro Kit according to the manufacturers’ protocols. Purified RNA was subjected to reverse transcription by using SuperScript III First-Strand Synthesis SuperMix for qRT-PCR. Quantification of gene expression was performed using the TaqMan Gene Expression Assays (ThermoFisher Scientific) and TaqMan Fast Advanced Master Mix. Data were collected by using QuantStudio 7 Flex Real-Time PCR System and analyzed by Excel and Prism. Results were normalized to YWHAH expression. Primer/probe used for qPCR are listed in the Table S2.

### Vector construction

Retrovirus plasmid encoding whole open reading frame (ORF) of human EGFR with C797S mutation was generated from pBabe_hEGFR by site-directed mutagenesis-like method. Three primer sets amplified either former-part EGFR (fragment1), latter-part EGFR (fragment2), or retrovirus backbone (fragment3), where common homology sequence between fragment1 and fragment2 contains C797S. Three fragments were ligated by using In-Fusion™ HD Cloning Plus.

Lentivirus plasmid encoding bicistronic TCRA and TCRB were generated as described previously^44^. Codon-optimized full-length TCR chains and truncated human CD19 (thCD19) marker gene, separated by a Furin-recognition and a self-cleaving 2A sequences, were synthesized (gBlocks). Using In-Fusion™ HD Cloning Plus, synthesized gene fragments were cloned into a third-generation lentiviral transfer vector plasmid, the backbone of which was prepared from CS_EF1a_iC9_mCherry treated by HindIII-HF and XbaI^45^.

Lentivirus plasmid encoding single-chained peptide-B2M-HLA trimer were generated by replacing peptide part with CS9.3 or CS11.6 using In-Fusion™ HD Cloning Plus. Original lentivirus plasmid was provided by Dr. David Baltimore.

### Virus vector production

Retroviral vector was produced by using GP2-293 system. A 10 cm culture dish was first treated by 0.01% Poly-L-Lysine (PLL) coating solution at room temperature for 1 hour. GP2-293 cells (2 × 10^5^ cells) were plated in the PLL-coated culture dish and cultured for 24 hours in D10 medium (DMEM + 10% FBS + 1% GlutaMax). Next, 10 µg of retroviral vector and 7.5 µg of pVSV-G were transfected into the cells using CalPhos™ Mammalian Transfection Kit. After 12 hours incubation, the medium was exchanged with fresh D10 medium and cultured for another 48 hours. The supernatant was collected and filtrated through 0.45 µm filter.

Lentiviral vector cloned with CS9.3- or CS11.6-recognizing TCRs were produced by using LV-MAX™ Lentiviral Production System following a manufacturer’s instruction. In brief, a high-density cultured suspension Viral Production Cells (4 × 10^7^ cells in 10 mL) were transfected with 10 µg of lentivirus plasmid, 7.5 µg of pMDLg/pRRE, and 7.5 µg of pCMV_VSV-G_RSV_Rev, and 0.4 mL of LV-MAX™ Enhancer added 5 hours after the transfection. At 48-55 hours post-transfection, culture medium was collected, centrifuged to exclude cells, filtrated through 0.45 µm filter, and quality confirmed by Lenti-X™ GoStix™ Plus.

### ProImmune REVEAL assay

Candidate peptide-HLA combination was selected by using the following cutoffs: BIMAS score, >1.000; IDEB rank, <10.0; or NetMHC, <10.0. Peptide-HLA binding affinity and stability assays were performed by high-throughput ProImmune REVEAL assays (ProImmune). Epitopes that bind HLA cause conformational changes and it was recognized by using a conformation-specific labeled antibody to provide a binding readout by ELISA. The REVEAL binding affinity score for each peptide-HLA complex is calculated by comparison with binding of the relevant positive control. Scores of more than 46% of control binding indicated significant or strong binders. Peptide-HLA combinations passed the binding affinity assay were then validated their binding stability of the resulting peptide-HLA complexes by using a quick check off-rate analysis. At 0, 2, and 24 hours samples of dissociating peptide-HLA complexes were taken from 37 °C also using conformational ELISA. One-phase dissociation curve, y = a*Exp(-k*x), was used where applicable to calculate the off-rates of the peptides. Half-life (τ1/2) values were calculated using the rate constant (k) in the following equation: τ1/2 = ln 2/k.

### Quantification and statistical analysis

Animal studies were performed without blinding and animals which couldn’t bear tumor were excluded from the analysis. No calculation for sample size estimation for animal studies. All assays were repeated at least three times. Two-tailed unpaired t test were performed using the GraphPad Prism program to determine statistical significance between groups. Error bars indicate standard deviation (SD) or standard error of the mean (SEM). P values less than 0.05 were considered to be statistically significant.

## ACKNOWLEDGEMENTS

We thank Miki Ando and Motoo Watanabe (The University of Tokyo) for helpful discussions; Naoto Hirano (University of Toronto) for kindly providing K562 aAPC; Gay Crooks (University of California Los Angeles) for kindly providing MS5-hDLL1 cell line; Mayumi Ono (Kyushu University) for kindly providing 11-18 cell line; and Peter Steinberger (Medical University of Vienna) for kindly providing Jurkat TPR cell line. We thank Hyam Levitsky (Johns Hopkins University) for helpful discussions and for critical reading of the manuscript.

The project was supported in part by the California Institute for Regenerative Medicine (LA1-06917 and DISC2-08874), Lung Cancer Research Foundation (SU ID: SPO 125372). K.N. was supported by a research fellowship from Japan Society for the Promotion of Science (JSPS). T.N. was supported by Weston Havens Foundation, and a research fellowship from Japan Society for the Promotion of Science (JSPS). J.L.F. was supported by the National Defense Science and Engineering Graduate (NDSEG) and Stanford Honorary Bio-X Fellowships. K.M.L. was supported by the NIH Director’s Early Independence Award (DP5OD024558) and is a Pew Scholar and The Anthony DiGenova Endowed Faculty Scholar.

## AUTHOR CONTRIBUTIONS

Conceptualization, T.N. and H.N.; Methodology, T.N., J.L.F., and K.M.L.; Investigation, T.N., K.N., J.V., L.A., J.L.F., T.I., Y.N., B.S., and H.X.; Resources, M.N. and J.B.S.; Writing – Original Draft, T.N., K.N., A.C.W., K.M.L., and H.N.; Supervision, K.M.L., J.B.S., and H.N.; Funding Acquisition, T.N., K.M.L., J.B.S., and H.N.

## SUPPLEMENTAL FIGURE LEGENDS

**Figure S1:**
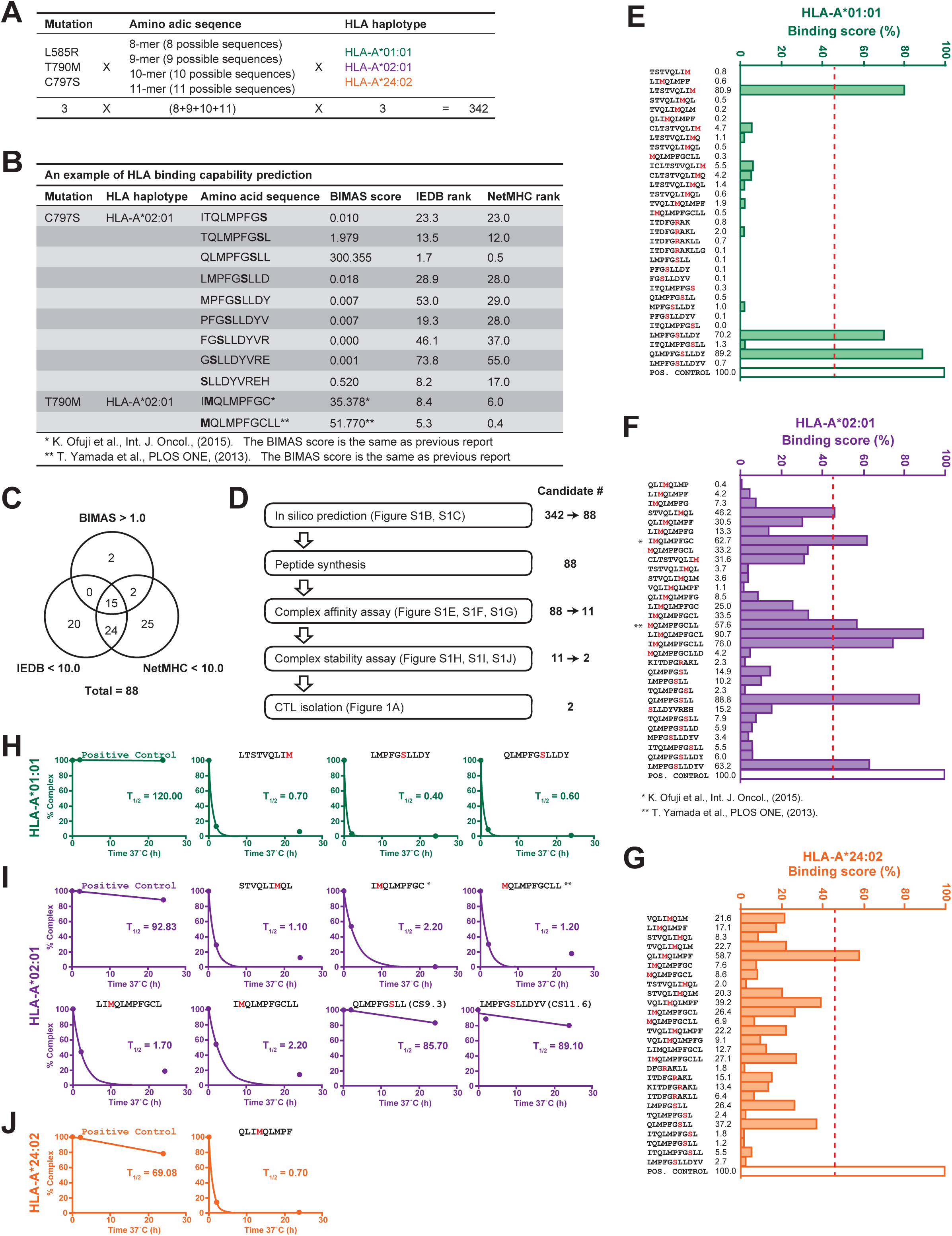
In silico and *in vitro* screening of EGFR neoantigen candidates. (A) Initial candidate size calculated by combination of EGFR mutations (L585R, T790M, C797S), amino acid sequences, and HLA haplotypes (HLA-A*01:01, HLA-A*02:01, HLA-A*24:02). All possible combinations = 342. (B) An example of the values after algorithms (BIMAS score, IEBD rank, NetMHC rank). Score: higher is more probable. Rank: lower is more probable. Two amino acid sequences and BIMAS scores on the bottom are immunogenic peptides derived from T790M mutation, described in previous reports. (C) The number of peptide sequences which met at least one of the selection criteria. (D) Workflow for *in vitro* screening to narrow down the number of neoantigen candidates. (E-G) *In vitro* binding of mutant EGFR-derived peptides to HLA A*01:01 (E), HLA-A*02:01 (F), or HLA A*24:02 (G). Each column shows the synthesized candidate peptide, 8 to 11 mer peptides with one amino acid substitution, and peptide-HLA binding score by the ProImmune REVEAL^®^ MHC-Peptide Binding Assays. Pos. control: control peptide with high affinity for HLA A*01:01, HLA-A*02:01, or HLA A*24:02 (set as 100% binging). Peptides with a binding score above 46% (dashed line) are considered positives. (H-J) *In vitro* assessment of long-term stability of peptide-HLA complex formation. For each peptide, the binding score at 0 hr, 2 hrs, and 24 hrs was measured.

**Figure S2:**
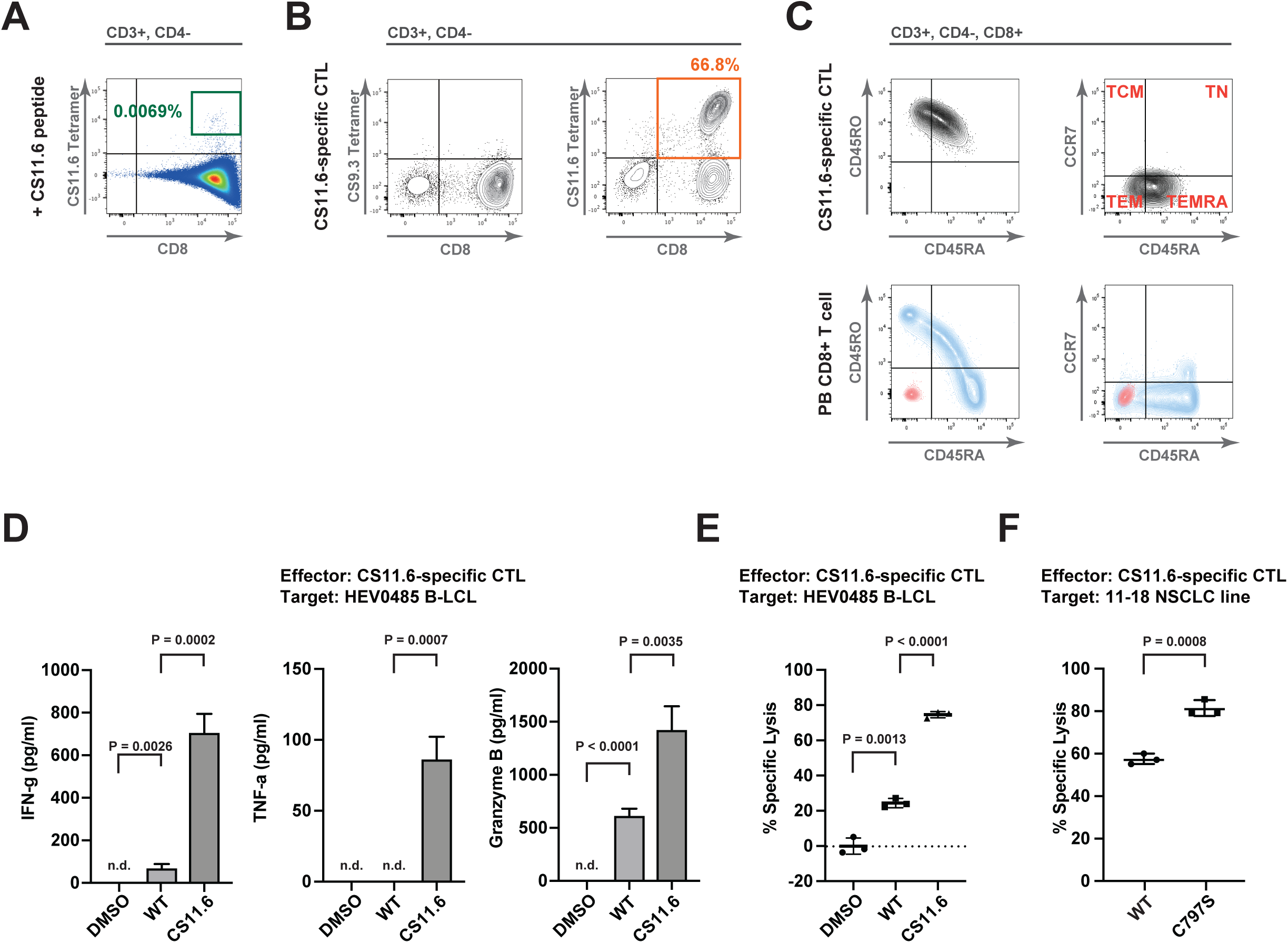
*In vitro* assessment of immunogenicity of EGFR neoantigens. (A) Flow cytometric analysis of the CD8^+^ T cells stimulated by peptide-pulsed autologous mDCs, 14 days after stimulation. Parent gate is indicated above each panel. (B) Flow cytometric analysis of purified and expanded CTL clones with cognate and irrelevant tetramers. Parent gate is indicated above each panel. (C) Flow cytometric analysis of memory phenotype in a CS11.6-specific CTL clone. Human peripheral blood CD8^+^ T cells (PB CD8+ T cell; bottom row) are shown as positive control (stained with the same antibody panel; blue) and negative control (unstained; red). Parent gate is indicated above each panel. (D) Cytometric Beads Array (CBA) analyses for IFN-γ, TNF-α, and Granzyme B secretion. CTL:HEV0485 = 10:1 (E:T ratio), 10 µg/ml peptide. (E) Antigen-specific cytotoxicity of CS11.6-specific CTL clone to HEV0485 B-LCLs. CTL:HEV0485 = 10:1 (E:T ratio), 10 µg/ml peptide. (F) Antigen-specific cytotoxicity of CS11.6-specific CTL clone to 11-18 NSCLC cell line. CTL:11-18 = 20:1 (E:T ratio). For (A-C), Data are representative of at least three independent experiments. For (D-F), Error bars = SD. Shown data are the representative in three times repeated experiments with technical triplicates in each experiment. P values are accorded to unpaired Student’s t test.

**Figure S3:**
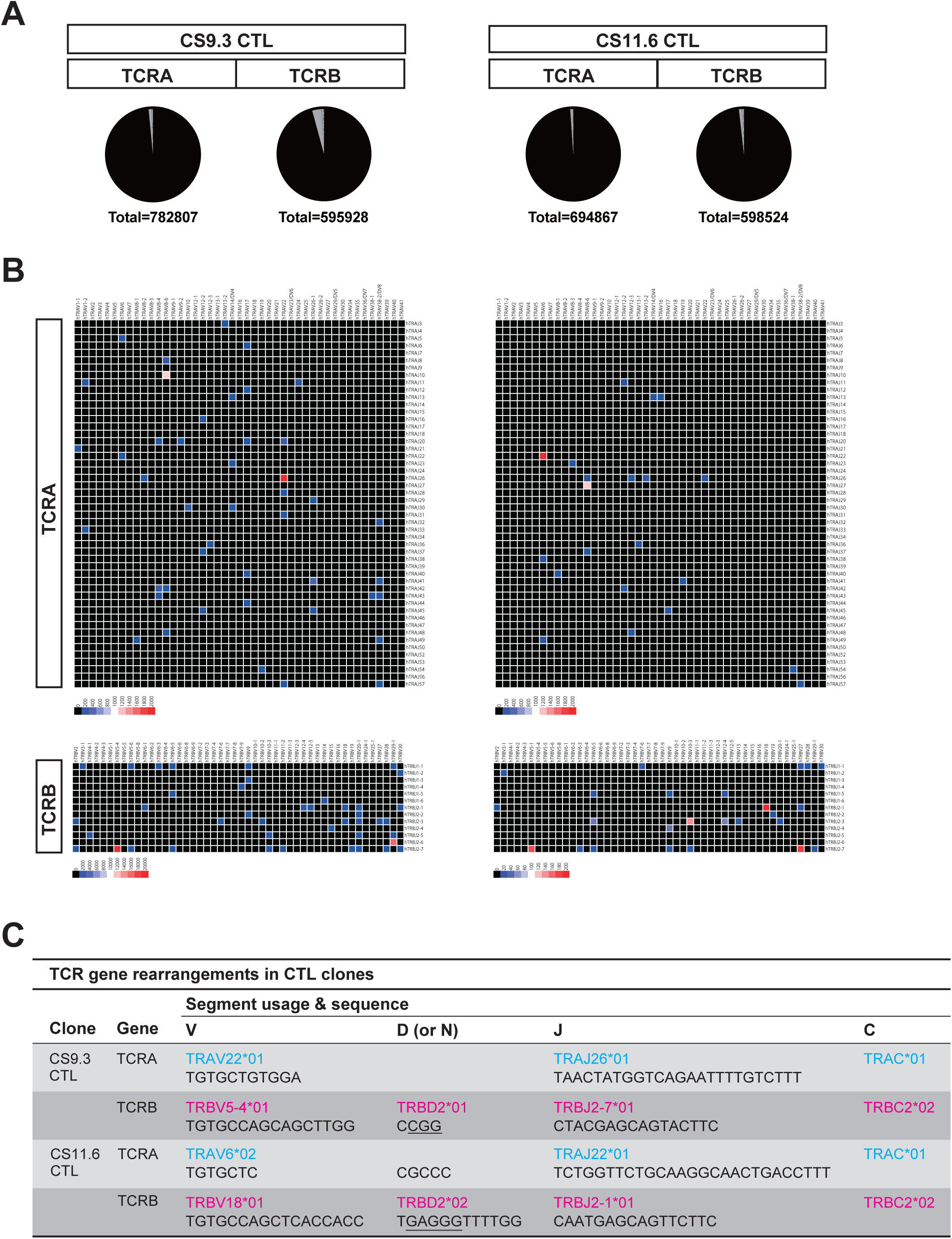
Identification of CS9.3- or CS11.6-specific TCRs. (A) Repertoire distribution of TCRA and TCRB in each CTL clone. Total read numbers for each TCR gene are indicated. (B) Heat map of the segment usage for TCRA and TCRB. (C) TCR gene rearrangements in CS9.3- or CS11.6-specific CTL clones. V, D, and J segment usages were identified by comparison to the ImMunoGeneTics (IMGT) database (http://www.imgt.org) and by using an on-line tool (IMGT/V-Quest)^46^. Gene-segment nomenclature follows IMGT usage.

**Figure S4:**
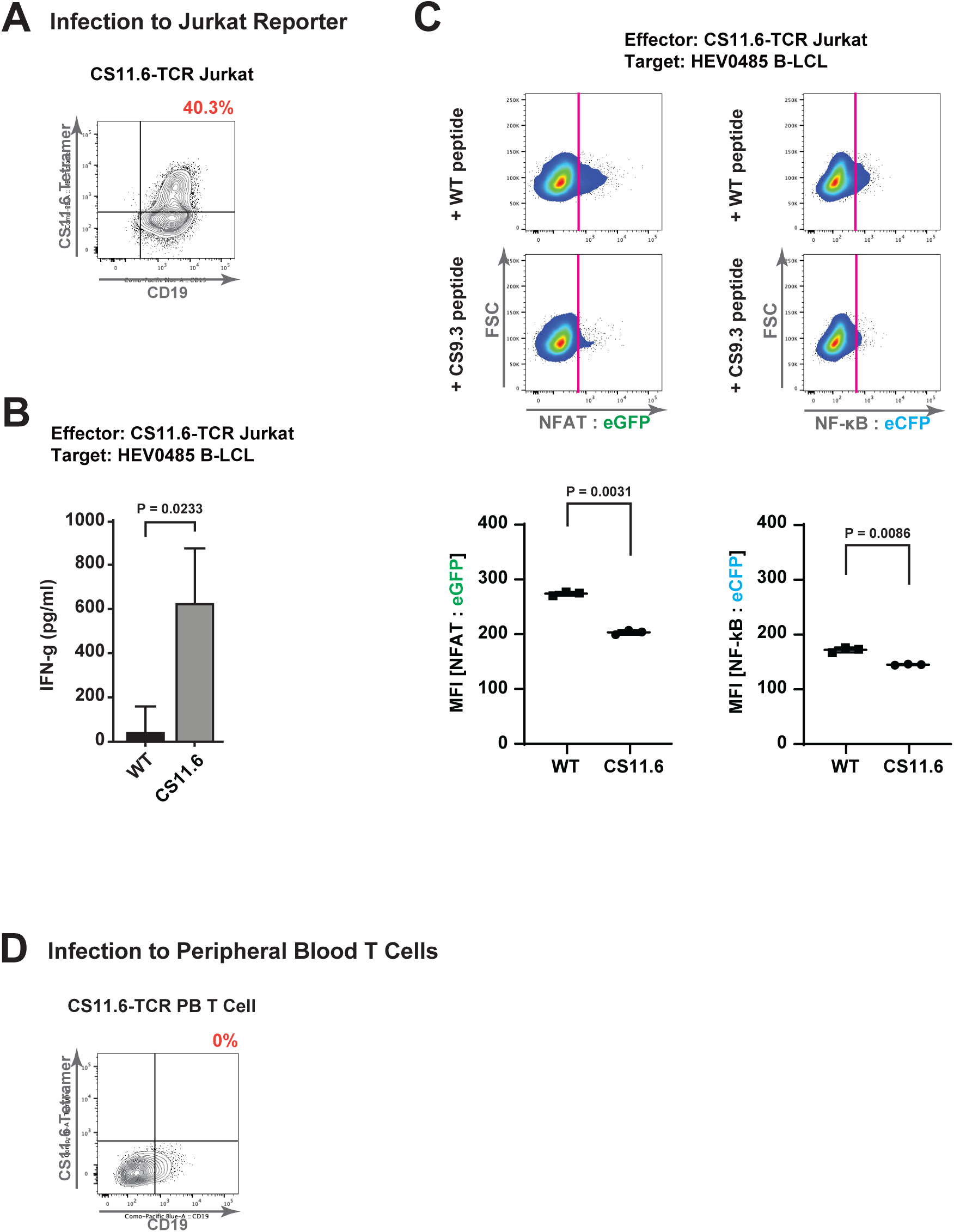
Functional evaluation of the TCR-transferred Jurkat reporter. (A) Flow cytometric analysis for tetramer recognition by using Jurkat reporter cells infected by lentivirus encoding CS11.6-specific TCRs. (B) Activation and secretion of IFN-γ in CS11.6-TCR transferred Jurkat reporter cells. Jurkat:HEV0485 = 10:1 (E:T ratio), 10 µg/ml peptide. (C) Flow cytometric analysis of CS11.6-TCR transferred Jurkat reporter cells. Jurkat:HEV0485 = 10:1 (E:T ratio), 10 µg/ml peptide. Bottom panels show the mean of fluorescence intensity (MFI) of reporters. (D) Flow cytometric analysis for tetramer recognition by using peripheral blood CD8^+^ CTLs infected by lentivirus encoding CS11.6-specific TCRs. For (B, and C), Error bars = SD. Shown data are the representative in three times repeated experiments with technical triplicates in each experiment. P values are accorded to unpaired Student’s t test.

**Figure S5:**
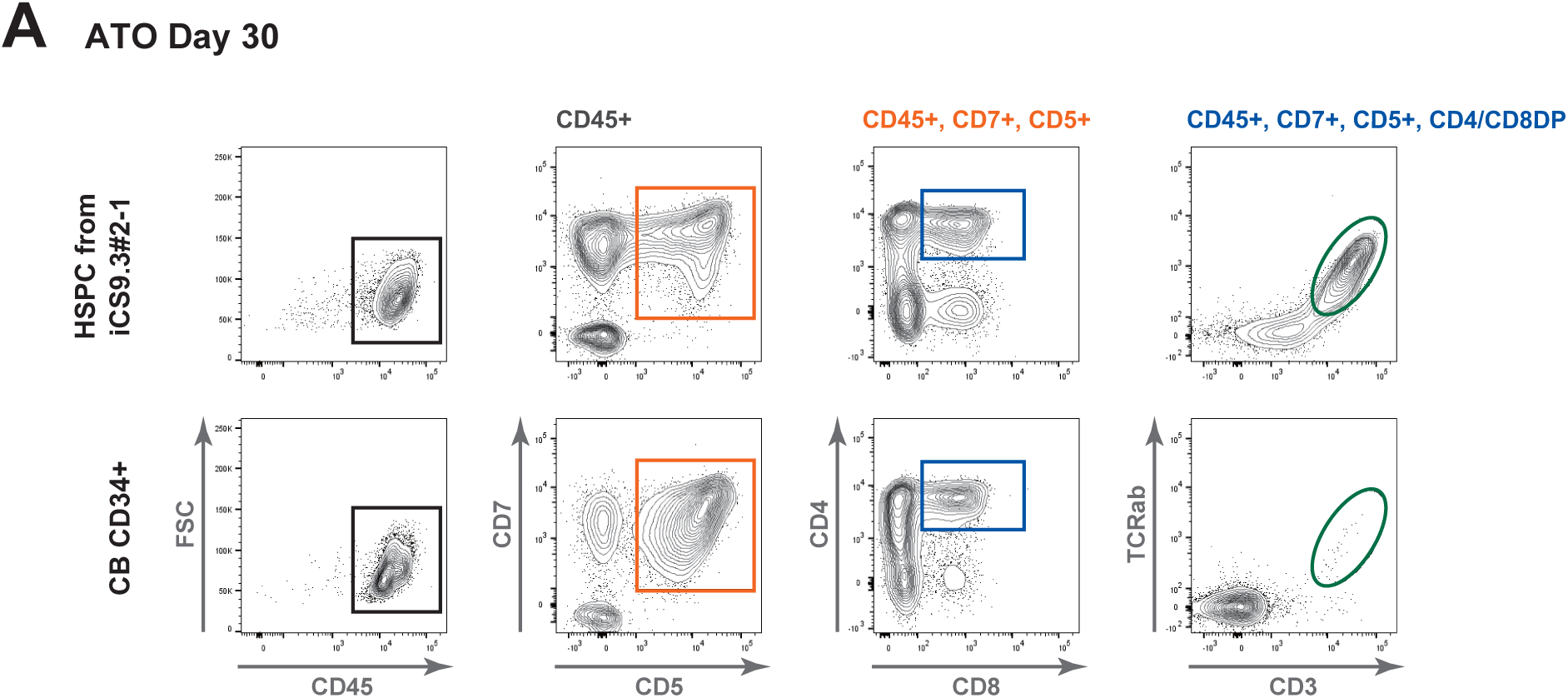
T-lineage specification by ATO culture. (A) Analysis of T-lineage specification by ATO culture (representative of three independent experiments) from CS9.3-specific T-iPSCs. CD34^+^ HSPCs purified from umbilical cord blood was differentiated in parallel as positive control for T-lineage specification by ATO culture. Subsequent parent gates are shown above each panel.

**Figure S6:**
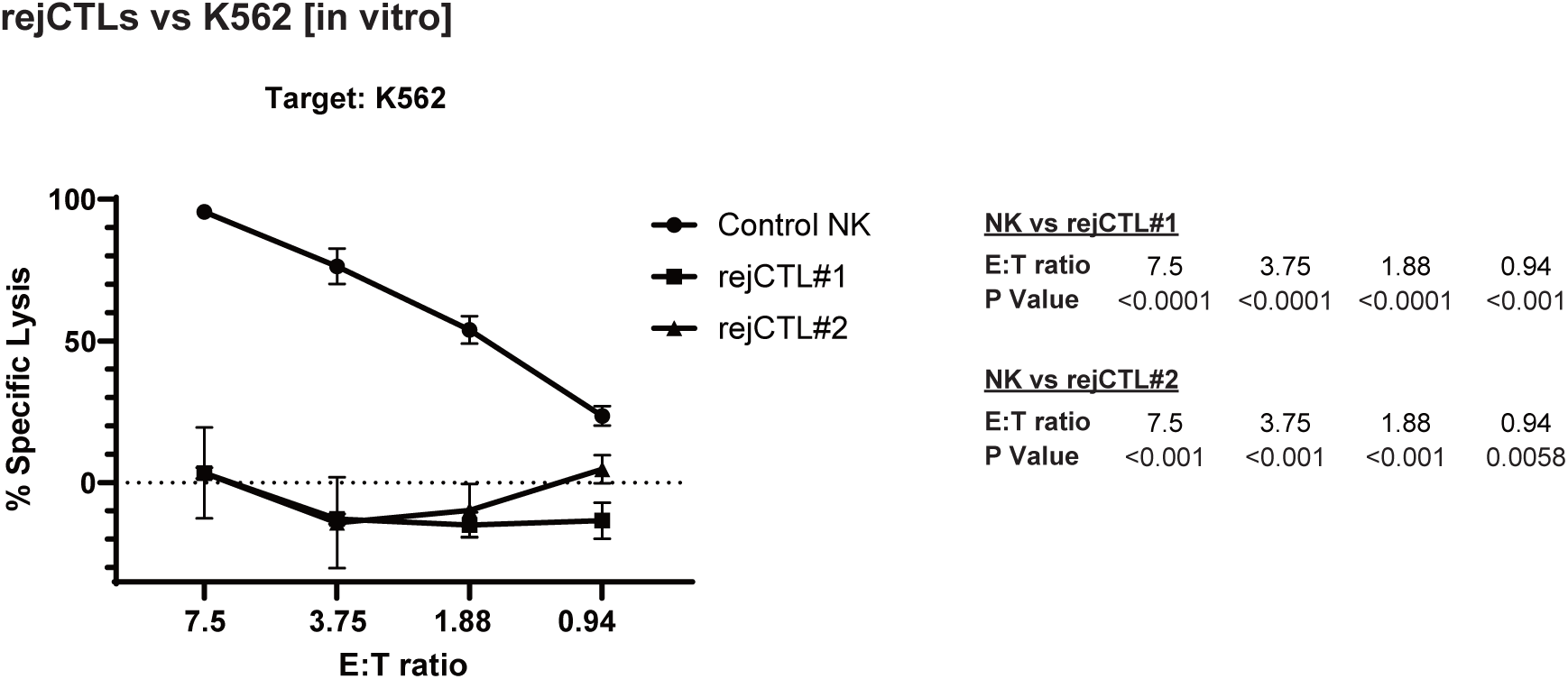
Assessments of lacking NK cell-like activity in iCTLs. Lack of NK cell-like cytotoxicity of iCTL clones to K562. Effector T cell:Target = 7.5:1, 3.75:1, 1.875:1, or 0.9375:1 (E:T ratio). Error bars = SD. Shown data are the representative in three times repeated experiments with technical triplicates in each experiment. P values are accorded to unpaired Student’s t test.

## KEY RESOURCES TABLE

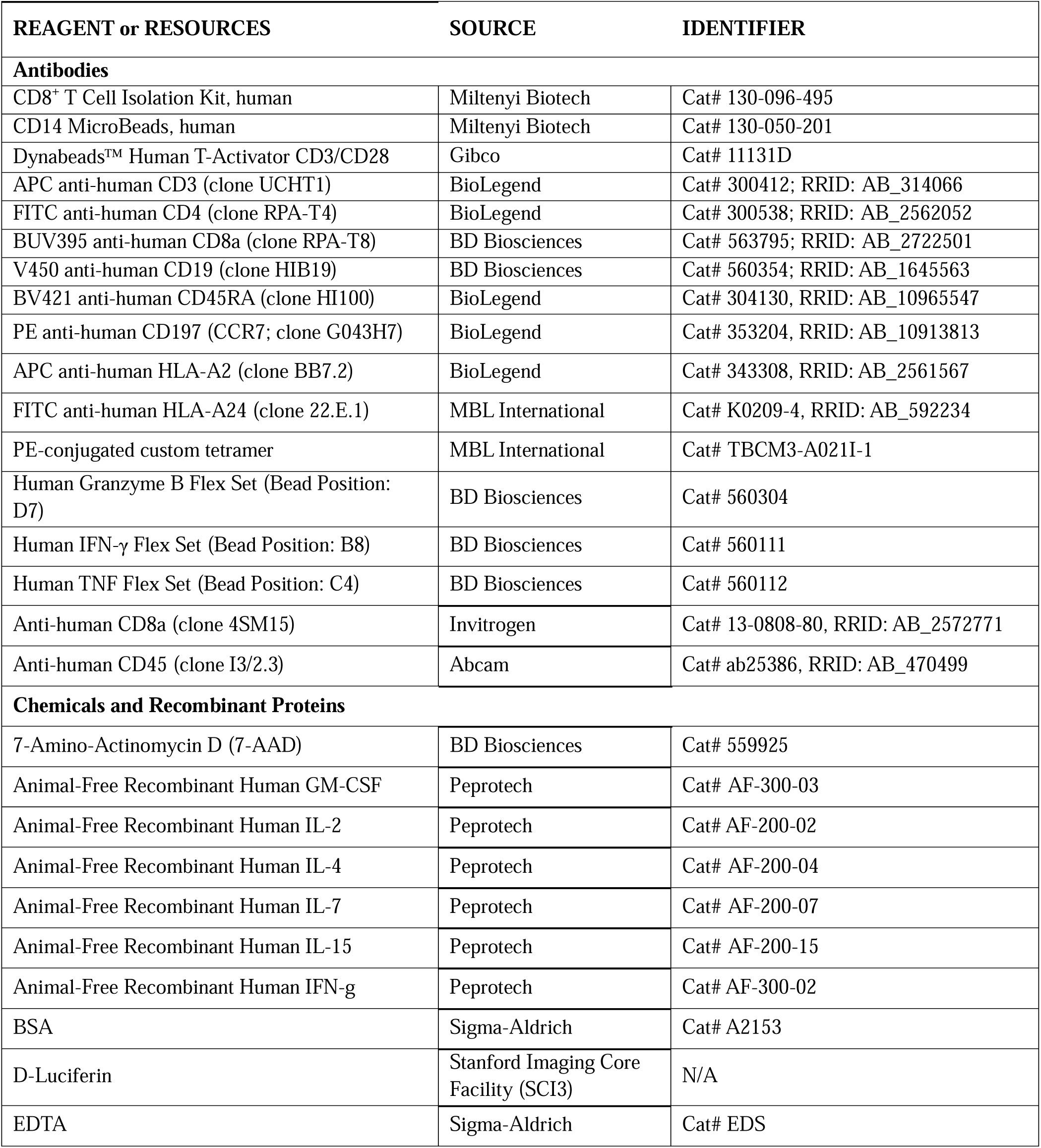
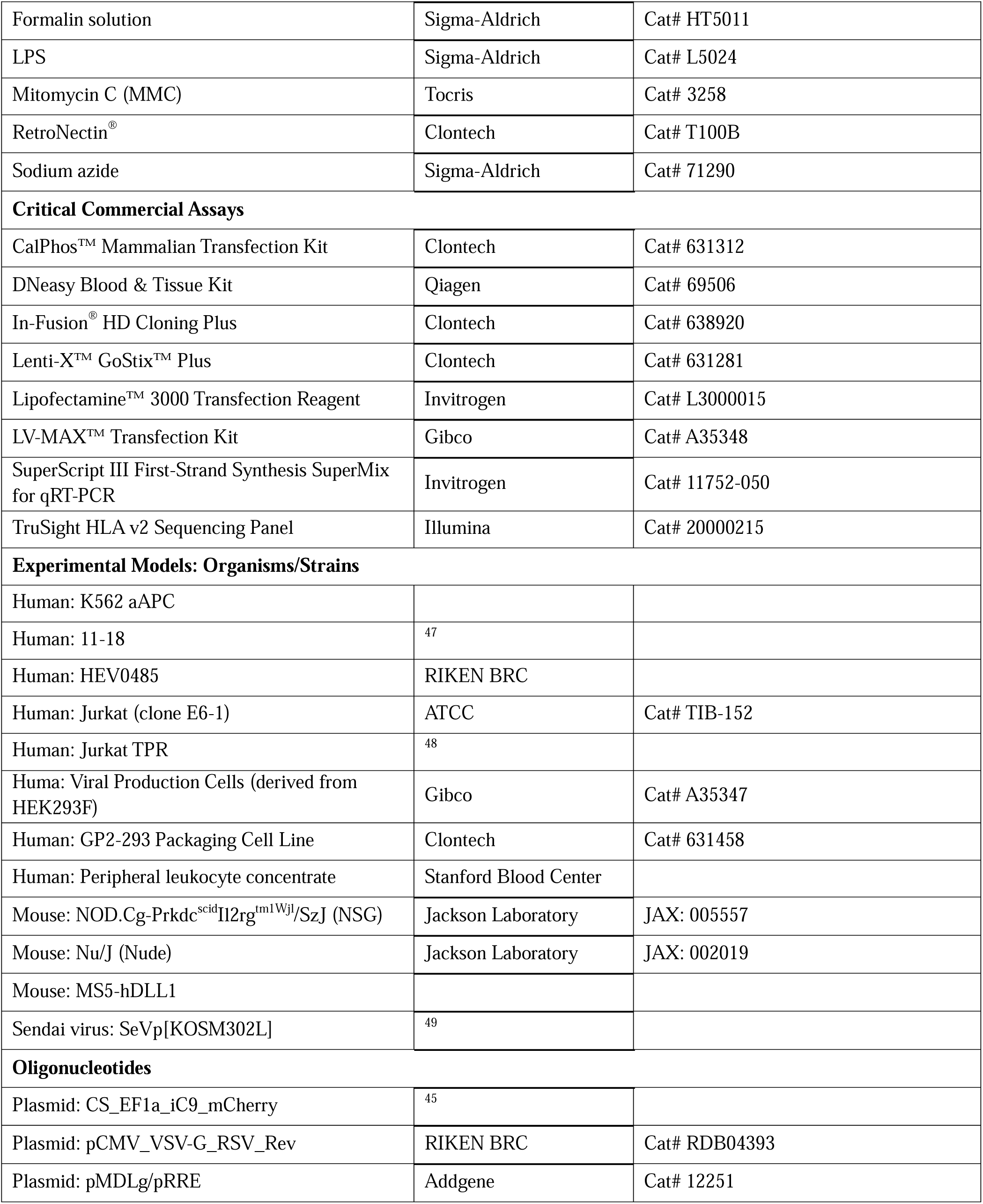
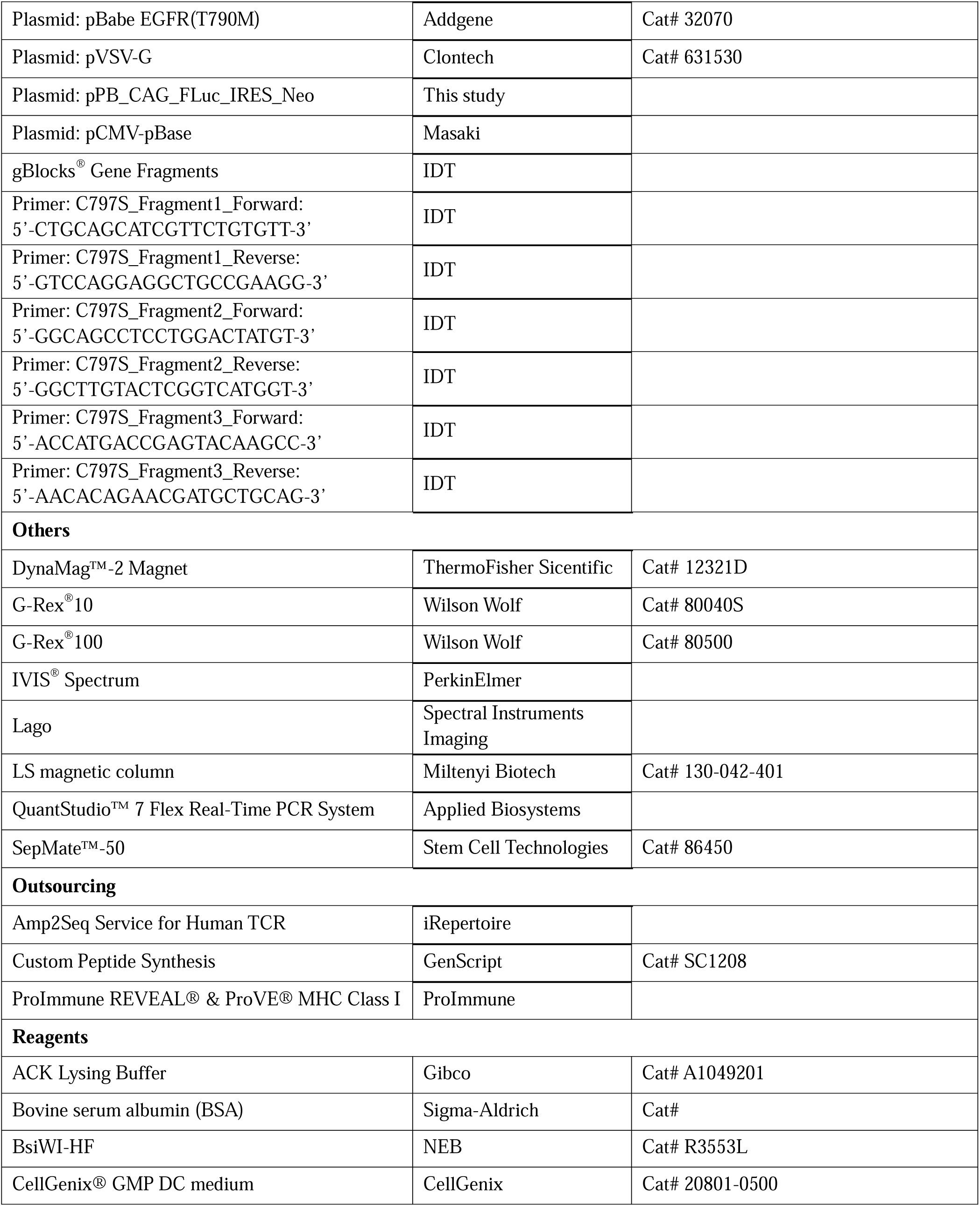
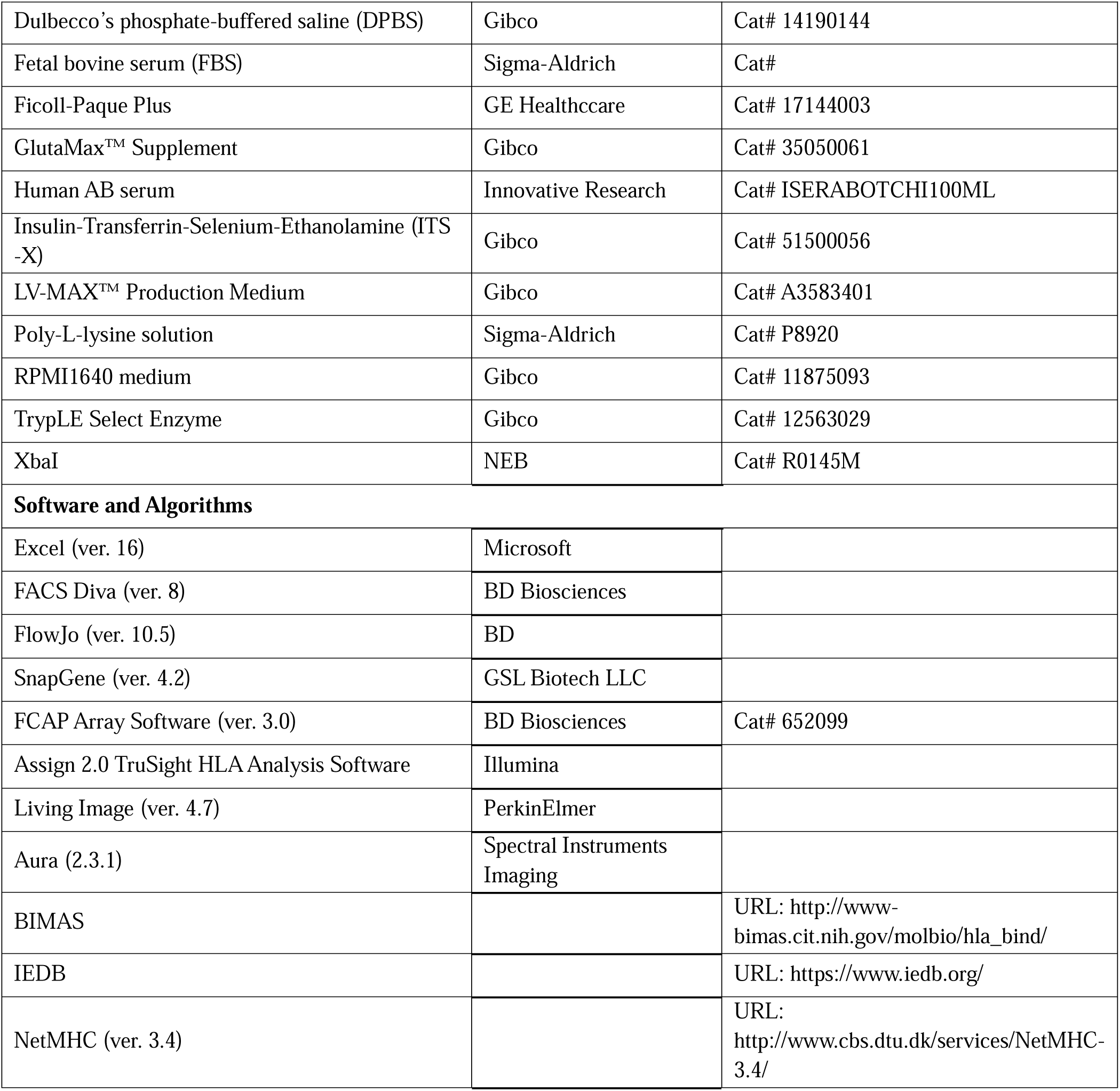

## REFERENCES

1. Yarchoan M, Johnson BA, 3rd, Lutz ER, Laheru DA, Jaffee EM. Targeting neoantigens to augment antitumour immunity. Nat Rev Cancer. Aug 24 2017;17(9):569. doi:10.1038/nrc.2017.74

2. Lim WA, June CH. The Principles of Engineering Immune Cells to Treat Cancer. Cell. Feb 9 2017;168(4):724–740. doi:10.1016/j.cell.2017.01.016

3. Tran E, Ahmadzadeh M, Lu YC, et al. Immunogenicity of somatic mutations in human gastrointestinal cancers. Science. Dec 11 2015;350(6266):1387–90. doi:10.1126/science.aad1253

4. Tran E, Robbins PF, Lu YC, et al. T-Cell Transfer Therapy Targeting Mutant KRAS in Cancer. N Engl J Med. Dec 8 2016;375(23):2255–2262. doi:10.1056/NEJMoa1609279

5. Themeli M, Kloss CC, Ciriello G, et al. Generation of tumor-targeted human T lymphocytes from induced pluripotent stem cells for cancer therapy. Nat Biotechnol. Oct 2013;31(10):928–33. doi:10.1038/nbt.2678

6. Ando M, Ando J, Yamazaki S, et al. Long-term eradication of extranodal natural killer/T-cell lymphoma, nasal type, by induced pluripotent stem cell-derived Epstein-Barr virus-specific rejuvenated T cells. Haematologica. Mar 2020;105(3):796–807. doi:10.3324/haematol.2019.223511

7. Maeda T, Nagano S, Ichise H, et al. Regeneration of CD8alphabeta T Cells from T-cell-Derived iPSC Imparts Potent Tumor Antigen-Specific Cytotoxicity. Cancer Res. Dec 1 2016;76(23):6839–6850. doi:10.1158/0008-5472.CAN-16-1149

8. Minagawa A, Yoshikawa T, Yasukawa M, et al. Enhancing T Cell Receptor Stability in Rejuvenated iPSC-Derived T Cells Improves Their Use in Cancer Immunotherapy. Cell Stem Cell. Dec 6 2018;23(6):850–858 e4. doi:10.1016/j.stem.2018.10.005

9. Nishimura T, Kaneko S, Kawana-Tachikawa A, et al. Generation of rejuvenated antigen-specific T cells by reprogramming to pluripotency and redifferentiation. Cell Stem Cell. Jan 3 2013;12(1):114–26. doi:10.1016/j.stem.2012.11.002

10. Themeli M, Riviere I, Sadelain M. New cell sources for T cell engineering and adoptive immunotherapy. Cell Stem Cell. Apr 2 2015;16(4):357–66. doi:10.1016/j.stem.2015.03.011

11. Reck M, Heigener DF, Mok T, Soria JC, Rabe KF. Management of non-small-cell lung cancer: recent developments. Lancet. Aug 24 2013;382(9893):709–19. doi:10.1016/S0140-6736(13)61502-0

12. Siegel RL, Miller KD, Jemal A. Cancer statistics, 2018. CA Cancer J Clin. Jan 2018;68(1):7–30. doi:10.3322/caac.21442

13. Zappa C, Mousa SA. Non-small cell lung cancer: current treatment and future advances. Transl Lung Cancer Res. Jun 2016;5(3):288–300. doi:10.21037/tlcr.2016.06.07

14. Yang Z, Hackshaw A, Feng Q, et al. Comparison of gefitinib, erlotinib and afatinib in non-small cell lung cancer: A meta-analysis. Int J Cancer. Jun 15 2017;140(12):2805–2819. doi:10.1002/ijc.30691

15. Singh J, Petter RC, Baillie TA, Whitty A. The resurgence of covalent drugs. Nat Rev Drug Discov. Apr 2011;10(4):307–17. doi:10.1038/nrd3410

16. Thress KS, Paweletz CP, Felip E, et al. Acquired EGFR C797S mutation mediates resistance to AZD9291 in non-small cell lung cancer harboring EGFR T790M. Nat Med. Jun 2015;21(6):560–2. doi:10.1038/nm.3854

17. Trolle T, McMurtrey CP, Sidney J, et al. The Length Distribution of Class I-Restricted T Cell Epitopes Is Determined by Both Peptide Supply and MHC Allele-Specific Binding Preference. J Immunol. Feb 15 2016;196(4):1480–7. doi:10.4049/jimmunol.1501721

18. Mei S, Li F, Leier A, et al. A comprehensive review and performance evaluation of bioinformatics tools for HLA class I peptide-binding prediction. Brief Bioinform. Jun 14 2019:bbz051. doi:10.1093/bib/bbz051

19. Harndahl M, Rasmussen M, Roder G, et al. Peptide-MHC class I stability is a better predictor than peptide affinity of CTL immunogenicity. Eur J Immunol. Jun 2012;42(6):1405–16. doi:10.1002/eji.201141774

20. Paul S, Weiskopf D, Angelo MA, Sidney J, Peters B, Sette A. HLA class I alleles are associated with peptide-binding repertoires of different size, affinity, and immunogenicity. J Immunol. Dec 15 2013;191(12):5831–9. doi:10.4049/jimmunol.1302101

21. Wolfl M, Greenberg PD. Antigen-specific activation and cytokine-facilitated expansion of naive, human CD8+ T cells. Nat Protoc. Apr 2014;9(4):950–66. doi:10.1038/nprot.2014.064

22. Fu X, Tao L, Rivera A, et al. A simple and sensitive method for measuring tumor-specific T cell cytotoxicity. PLoS One. Jul 29 2010;5(7):e11867. doi:10.1371/journal.pone.0011867

23. Kanda R, Kawahara A, Watari K, et al. Erlotinib resistance in lung cancer cells mediated by integrin beta1/Src/Akt-driven bypass signaling. Cancer Res. Oct 15 2013;73(20):6243–53. doi:10.1158/0008-5472.CAN-12-4502

24. Hayashi K, Ishizuka S, Yokoyama C, Hatae T. Attenuation of interferon-gamma mRNA expression in activated Jurkat T cells by exogenous zinc via down-regulation of the calcium-independent PKC-AP-1 signaling pathway. Life Sci. Jul 4 2008;83(1-2):6–11. doi:10.1016/j.lfs.2008.04.022

25. Vizcardo R, Klemen ND, Islam SMR, et al. Generation of Tumor Antigen-Specific iPSC-Derived Thymic Emigrants Using a 3D Thymic Culture System. Cell Rep. Mar 20 2018;22(12):3175–3190. doi:10.1016/j.celrep.2018.02.087

26. Fowler JL, Zheng SL, Nguyen A, et al. Lineage-tracing hematopoietic stem cell origins in vivo to efficiently make human HLF+ HOXA+ hematopoietic progenitors from pluripotent stem cells. Dev Cell. May 06 2024;59(9):1110–1131.e22. doi:10.1016/j.devcel.2024.03.003

27. Montel-Hagen A, Seet CS, Li S, et al. Organoid-Induced Differentiation of Conventional T Cells from Human Pluripotent Stem Cells. Cell Stem Cell. Mar 7 2019;24(3):376–389 e8. doi:10.1016/j.stem.2018.12.011

28. Seet CS, He C, Bethune MT, et al. Generation of mature T cells from human hematopoietic stem and progenitor cells in artificial thymic organoids. Nat Methods. May 2017;14(5):521–530. doi:10.1038/nmeth.4237

29. Deuse T, Hu X, Gravina A, et al. Hypoimmunogenic derivatives of induced pluripotent stem cells evade immune rejection in fully immunocompetent allogeneic recipients. Nat Biotechnol. 03 2019;37(3):252–258. doi:10.1038/s41587-019-0016-3

30. Wang B, Iriguchi S, Waseda M, et al. Generation of hypoimmunogenic T cells from genetically engineered allogeneic human induced pluripotent stem cells. Nat Biomed Eng. May 2021;5(5):429–440. doi:10.1038/s41551-021-00730-z

31. Furukawa Y, Ishii M, Ando J, et al. iPSC-derived hypoimmunogenic tissue resident memory T cells mediate robust anti-tumor activity against cervical cancer. Cell Rep Med. Dec 19 2023;4(12):101327. doi:10.1016/j.xcrm.2023.101327

32. Lo W, Parkhurst M, Robbins PF, et al. Immunologic Recognition of a Shared p53 Mutated Neoantigen in a Patient with Metastatic Colorectal Cancer. Cancer Immunol Res. Apr 2019;7(4):534–543. doi:10.1158/2326-6066.CIR-18-0686

33. Akazawa Y, Saito Y, Yoshikawa T, et al. Efficacy of immunotherapy targeting the neoantigen derived from epidermal growth factor receptor T790M/C797S mutation in non-small cell lung cancer. Cancer Sci. Aug 2020;111(8):2736–2746. doi:10.1111/cas.14451

34. Sidney J, Southwood S, Mann DL, Fernandez-Vina MA, Newman MJ, Sette A. Majority of peptides binding HLA-A*0201 with high affinity crossreact with other A2-supertype molecules. Hum Immunol. Nov 2001;62(11):1200–16.

35. Bentzen AK, Hadrup SR. T-cell-receptor cross-recognition and strategies to select safe T-cell receptors for clinical translation. Immunooncol Technol. Sep 2019;2:1–10. doi:10.1016/j.iotech.2019.06.003

36. Dudley ME, Wunderlich JR, Robbins PF, et al. Cancer regression and autoimmunity in patients after clonal repopulation with antitumor lymphocytes. Science. Oct 25 2002;298(5594):850–4. doi:10.1126/science.1076514

37. June CH, Sadelain M. Chimeric Antigen Receptor Therapy. N Engl J Med. Jul 5 2018;379(1):64–73. doi:10.1056/NEJMra1706169

38. Shifrut E, Carnevale J, Tobin V, et al. Genome-wide CRISPR Screens in Primary Human T Cells Reveal Key Regulators of Immune Function. Cell. Dec 13 2018;175(7):1958–1971 e15. doi:10.1016/j.cell.2018.10.024

39. Gornalusse GG, Hirata RK, Funk SE, et al. HLA-E-expressing pluripotent stem cells escape allogeneic responses and lysis by NK cells. Nat Biotechnol. Aug 2017;35(8):765–772. doi:10.1038/nbt.3860

40. Tran E, Robbins PF, Rosenberg SA. ’Final common pathway’ of human cancer immunotherapy: targeting random somatic mutations. Nat Immunol. Feb 15 2017;18(3):255–262. doi:10.1038/ni.3682

41. Hu Z, Ott PA, Wu CJ. Towards personalized, tumour-specific, therapeutic vaccines for cancer. Nat Rev Immunol. Mar 2018;18(3):168–182. doi:10.1038/nri.2017.131

42. Kawana-Tachikawa A, Tomizawa M, Nunoya J, et al. An efficient and versatile mammalian viral vector system for major histocompatibility complex class I/peptide complexes. J Virol. Dec 2002;76(23):11982–8. doi:10.1128/jvi.76.23.11982-11988.2002

43. Miyazaki T, Isobe T, Nakatsuji N, Suemori H. Efficient Adhesion Culture of Human Pluripotent Stem Cells Using Laminin Fragments in an Uncoated Manner. Sci Rep. 01 2017;7:41165. doi:10.1038/srep41165

44. Legut M, Dolton G, Mian AA, Ottmann OG, Sewell AK. CRISPR-mediated TCR replacement generates superior anticancer transgenic T cells. Blood. Jan 18 2018;131(3):311–322. doi:10.1182/blood-2017-05-787598

45. Ando M, Nishimura T, Yamazaki S, et al. A Safeguard System for Induced Pluripotent Stem Cell-Derived Rejuvenated T Cell Therapy. Stem Cell Reports. Oct 13 2015;5(4):597–608. doi:10.1016/j.stemcr.2015.07.011

46. Lefranc MP. IMGT databases, web resources and tools for immunoglobulin and T cell receptor sequence analysis, http://imgt.cines.fr. Leukemia. Jan 2003;17(1):260–6. doi:10.1038/sj.leu.2402637

47. Kawahara A, Yamamoto C, Nakashima K, et al. Molecular diagnosis of activating EGFR mutations in non-small cell lung cancer using mutation-specific antibodies for immunohistochemical analysis. Clin Cancer Res. Jun 15 2010;16(12):3163–70. doi:10.1158/1078-0432.CCR-09-3239

48. Jutz S, Hennig A, Paster W, et al. A cellular platform for the evaluation of immune checkpoint molecules. Oncotarget. Sep 12 2017;8(39):64892–64906. doi:10.18632/oncotarget.17615

49. Nishimura K, Sano M, Ohtaka M, et al. Development of defective and persistent Sendai virus vector: a unique gene delivery/expression system ideal for cell reprogramming. J Biol Chem. Feb 11 2011;286(6):4760–71. doi:10.1074/jbc.M110.183780

